# BiLSTM-Powered Bilinear Attention for Protein–Ligand Prediction

**DOI:** 10.64898/2026.05.10.724184

**Authors:** Chih-Yang Cheng, Yi-An Chen, Feng-Yin Li, Suyong Re

**Affiliations:** Laboratory of In Silico Design, Artificial Intelligence Center for Health and Biomedical Research, National Institutes of Biomedical Innovation, Health and Nutrition, Osaka, Japan; Department of Chemistry, National Chung Hsing University, Taichung 402, Taiwan

**Keywords:** Protein-ligand interaction, Weakly supervised learning, Graph Convolutional Networks, Long-range sequence modeling, Interpretability

## Abstract

Rapid and accurate prediction of protein-ligand bindings is essential for drug discovery. While generative AI has driven rapid advancements in structure-based approaches, sequence-based methods remain significantly faster and more cost-effective. Here, we present a weakly supervised deep learning framework integrating graph convolutional networks (GCN) for molecular encoding and bidirectional long short-term memory (BiLSTM) for protein modeling. The latter represents long-range dependencies better than the widely used convolutional neural network (CNN). Leveraging a bilinear attention network (BAN), this model learns protein-ligand pairwise interactions without requiring three-dimensional structural supervision. By using the publicly available BindingDB dataset, the model was trained, solely on affinity labels, and successfully classified binder and non-binders with AUROC of 0.96 and an AUPRC of 0.95. The model generates interpretable attention maps that serve as a “GPS” to locate binding sites. Remarkably, despite the lack of structural training data, it can pinpoint key contact residues confirmed by crystal structures. Our method could function as a scalable filter for giga-scale libraries, allowing rapid screening of drug candidates with direct structural insights into the protein-ligand interface.

## 1 Introduction

Developing new drugs is a highly cost-intensive endeavor, often costing billions of dollars.^1,2^ While the integration of Artificial Intelligence (AI) is dramatically transforming the efficiency of virtual drug screening such as through data-driven representation learning,^3–5^ a fundamental challenge remains: how to quickly and accurately identify novel compounds from a chemical space estimated at over 10^60^ molecules.^6,7^ Consequently, there is an urgent need for methods that can effectively expand the search space without compromising the accuracy of protein–ligand interaction (PLI) predictions, from binding site identification and pose prediction to lead optimization.^8^

High-accuracy protein structure prediction models like AlphaFold have driven a paradigm shift in structure-based drug design.^9,10^ AlphaFold3, in particular, has expanded the scope of virtual screening by providing high-fidelity structures of not only proteins but also complex biomolecular assemblies. Complementarily, models like DrugCLIP^11^ leverage cross-modal alignment between chemical structures and binding pockets to streamline the identification of potential ligands. These advances have overcome traditional limitations due to the scarcity of experimental structures.^12,13^ However, their predictive accuracy remains to be verified, and the high computational cost of docking simulations still hinders large-scale screening.^14^

As an alternative to computationally expensive structural approaches, sequence-based methods have emerged as a high-throughput and robust frontier.^15,16^ These methods bypass the requirement for three-dimensional structures by treating protein-ligand interactions as a representation learning problem. Deep neural networks (including convolutional neural networks (CNNs) for local motif extraction) are employed to achieve performance levels that rival traditional structure-based drug design. Leveraging protein language models and molecular graphs to a graph-to-transformer cross-attention mechanism, a screening power is further enhanced.^17,18^ A notable advancement in this domain is DrugBAN,^4^ which utilizes a bilinear attention network (BAN)^24^ to capture fine-grained reciprocal interactions between drug and target representations, effectively identifying sub-structural binding patterns without explicit docking (weakly supervised).

Here, we propose a weakly supervised end-to-end deep learning method that shares the basic framework with DrugBAN. The fundamental difference from DrugBAN is that we deliberately employ a bidirectional long short-term memory (BiLSTM) architecture instead of CNN in order to capture the global sequence dependencies critical for identifying non-contiguous binding residues.^20–22^ Adapted from the DrugBAN framework, we integrate a hybrid representation (BiLSTM^20,21^ for protein and graph convolutional networks (GCN)^23^ for ligand) with a BAN which allows robust co-attention learning without the cost of full outer-product representations by employing low-rank bilinear pooling. Trained solely on affinity data using BindingDB datasets,^24^ our model not only yielded high predictive performance (AUROC of 0.96 and AUPRC of 0.95) but also successfully pinpointed key contact residues confirmed by crystal structures. The presented model offers a high-throughput pre-screening engine to complement intensive structural models, enabling a hierarchical paradigm for large scale drug screening.

## 2 Methods

### Overall Architecture

The overall architecture of the proposed framework is illustrated in Fig. 1. Our model integrates ligand 2D graphs (SMILES-derived nodes and edges) and protein amino acid sequences (in FASTA format). The hybrid encoding architecture (Fig. 1a) leverages a GCN^23^ for extracting molecular topology of ligands and a BiLSTM^20,21^ for encoding sequential dependency of proteins. A similar hybrid architecture, combining BiLSTM with temporal convolutional networks (TCN), has proven effective for predicting protein secondary structures.^25^ These encoded features are fused via a BAN (Fig. 1b),^24^ which captures the non-linear correlation between ligand substructures and protein residues through a pairwise interaction map. Unlike simple concatenation, the BAN’s attention weights reflect multiplicative protein-ligand interactions. This joint representation is then processed by a fully connected decoder to predict binding probability (Fig. 1c). The learned attention weights can be projected onto the 3D protein structure (Fig. 1d), providing mechanistic interpretability by localizing potential binding sites.

**Fig. 1.**
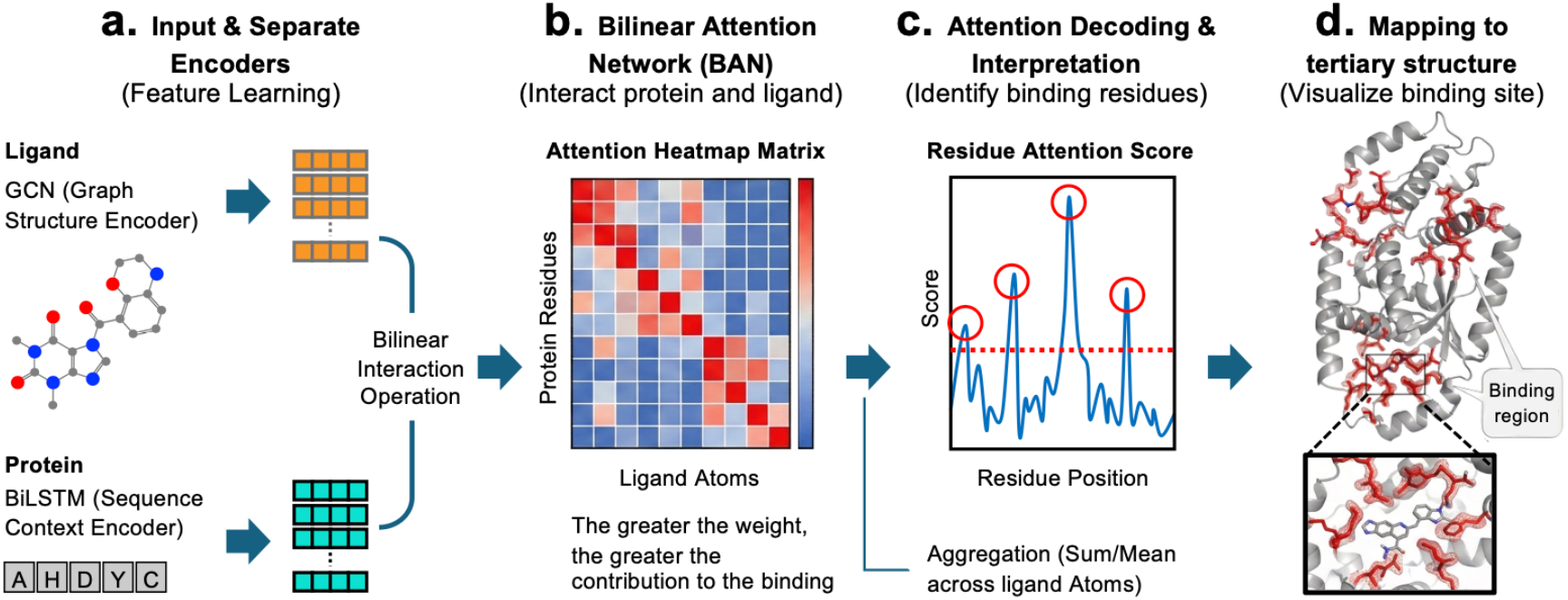
Overall architecture: a weakly supervised graph-sequence interaction framework. (a) The pipeline processes ligand molecular graphs and protein sequences through a hybrid encoder combining Graph Convolutional Networks (GCN) and Bidirectional LSTMs (BiLSTM). (b) Encoded features are integrated using a Bilinear Attention Network (BAN) and (c) then processed by a decoder to predict binding affinity. (d) Finally, the model generates interpretable attention maps that localize potential binding sites on the 3D structure, achieved solely from sequence data without coordinate supervision (weakly supervised).

### Datasets

A curated subset of the BindingDB dataset,^24^ specifically processed to minimize data bias, was used. This dataset comprises 49,199 drug-target interaction pairs, involving 14,643 unique ligands and 2,623 target proteins. The dataset was partitioned into training, validation, and test sets in a 7:1:2 ratio.

### Feature Representation and Encoding

#### Ligand Encoding with GCN

To preserve the topological structures of small molecules, we employed GCN as the ligand encoder.^23^ The ligand SMILES strings were first converted into 2D molecular graphs using RDKit.^26^ Each ligand molecule is represented as an undirected graph *G* = (*V, E*), where nodes *V* and edges *E* correspond to atoms and chemical bonds, respectively. To standardize the input feature space, we defined an initial node feature vector of dimension *D* = 78 for each atom, incorporating key physicochemical properties: atom type (one-hot encoded for C, N, O, etc.), node degree, formal charge, radical electrons, hybridization, aromaticity, hydrogens, and chirality. Three GCN layers were stacked to iteratively aggregate neighboring information, refining local structure representations. This graph-based approach explicitly captures chemical connectivity and geometry, which outperforms sequence-based ligand encoders (such as BiLSTM) by providing a more topologically faithful representation.^16^

#### Protein Encoding with BiLSTM

We strategically adopted the BiLSTM architecture to optimize the trade-off between representational capacity and computational efficiency. While CNNs excel at extracting local motifs, their fixed kernel sizes inherently limit their ability to capture long-range sequential dependencies. Such dependencies are fundamental to protein tertiary structure formation, where distant residues interact in 3D space.^15,22^ While BiLSTM involves higher computational cost, its recurrent mechanism effectively models these global contexts, surpassing the inherent limitations of CNN-based encoders.

### BAN: Bilinear Attention Network for Interaction Modeling

The BAN proposed by Kim et al.^19^ was used to model the fine-grained biochemical interactions between ligands and protein residues. Unlike conventional attention mechanisms that rely on simple feature concatenation, the BAN facilitates multiplicative interactions through a bilinear pooling scheme. This allows the model to capture complex, non-linear dependencies between the ligand and protein that are often missed by additive methods. Let 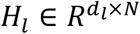 denote the encoded ligand representation, where *N* is the number of atoms and *d*_*l*_ is the GCN feature dimension (e.g., 128). Similarly, the protein representation is defined as 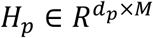, where *M* is the protein sequence length and *d*_*p*_ is the BiLSTM hidden dimension (e.g., 2 × hidden units for bidirectional context).

#### Bilinear Projection and Pooling

To align these heterogeneous feature spaces, we employ two learnable projection matrices, 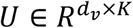 and 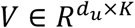, which map the ligand and protein features into a common *K*-dimensional joint semantic space. The bilinear attention map *A* ∈ *R*^*N*×*M*^ is computed using the Hadamard product (·) as follows:

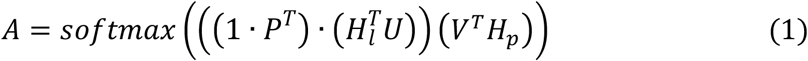

Where *P* ∈ *R*^*K*^ is a learnable weight vector, and 1 ∈ *R*^*N*^ is an all-ones vector. The term 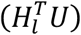 results in an *N* × *K* matrix, and 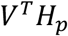 results in a *K* × *M* matrix. The final attention map *A* captures the pairwise interaction intensity between every ligand substructure and protein residue.^4,19^ The joint representation *f* is then obtained by pooling the weighted interaction features, enabling the model to focus on critical binding regions without prior structural supervision (weakly supervised).

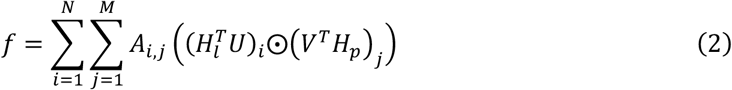

where 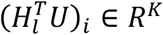 denotes the *i*-th row vector of the projected ligand features,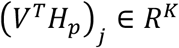 denotes the *j*-th column vector of the projected protein features, and ⨀ represents the Hadamard product (element-wise multiplication).

#### Prediction Head

We formulated the prediction task as a binary classification problem to distinguish active binders from non-binders. Consequently, the model’s final output is a probability score ŷ ∈ [0,1], representing the likelihood of a specific binding event. To ensure a rigorous decision boundary and mitigate noise in bioactivity data, we adopted a strict thresholding strategy: protein-ligand pairs with *IC*_*50*_ < 100 nM were labeled as positive, while those with *IC*_*50*_ > 10,000 nM were labeled as negative. Samples in the intermediate range (100 to 10,000 nM) were excluded to eliminate ambiguity. Crucially, this binary formulation does more than categorize interactions; it compels the BAN to learn the underlying geometric constraints of the binding interface. By optimizing for classification accuracy, the model spontaneously assigns higher attention weights to key residues, effectively generating an interpretable, residue-level importance map.

### Interpretation and Binding Site Localization

While being trained solely on binary affinity labels (*IC*_*50*_), a key advantage of our framework is its inherent interpretability, as inherited from the original DrugBAN.^4^ The BAN module spontaneously highlights residues critical for ligand binding through its learned attention weights.

#### Consensus Attention Aggregation

To minimize noise from each prediction and identify robust binding pockets, we employed a “one-target-multiple-ligands” aggregation strategy. That is, for a specific target protein consisting of *L* amino acid residues, we collected a set of *N* known active ligands {*d*_1_, *d*_2_, …, *d*_*N*_}. First, we obtained the raw bilinear attention map for each protein-ligand pair. Let *a*_*i,j*_ denote the attention weight assigned to the *j*-th amino acid residue by the *i*-th ligand. The consensus attention score *w*_*j*_ for residue *j* is obtained by averaging across all *N* ligands.

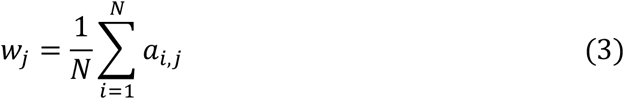

This aggregation filters out ligand-specific noise and amplifies the signal of conserved binding residues. To identify high-confidence residues within the target protein sequence, we standardized the attention weights using Z-score. The score *Z*_*j*_ for each residue *j* is calculated relative to the weight distribution across the entire protein sequence as follows:

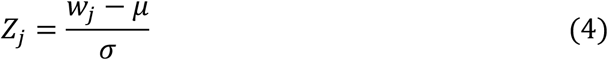

where *µ* and *σ* are the mean and standard deviation of the consensus weights (*w*_1_, …, *w*_*L*_) for the target protein. Residues with *Z*_*j*_ exceeding a specific threshold (corresponding to the top 75th percentile) were designated as candidate binding sites.

#### Residue Frequency Analysis

To analyze the biochemical properties of the predicted binding sites, we calculated the composition frequency of the identified residues. Let *R*_*pred*_ be the set of residues predicted as binding sites. The frequency *f* (*a*) for amino acid type *a* (e. g., Lys, Glu) is:

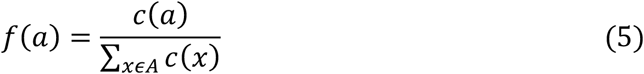

where *c*(*a*) is the count of amino acid type *a* within *R*_*pred*_, and *A* is the set of 20 standard amino acids. This profile enables us to verify whether the predicted pocket is enriched with binding-relevant physicochemical properties, such as hydrophobic or charged residues, relative to the background sequence. For structural validation, we mapped these high-Z-score residues onto 3D crystal structures from PDBbind^27^ and quantified their overlap with ground-truth pockets, defined as residues within 4.0 Å of the ligand.

#### Implementation and Training Details

The model was implemented using PyTorch.^28^ We optimized the network with the Adam optimizer at a learning rate of 5 × 10^56^ and a batch size of 64, employing binary cross-entropy as the loss function.^29^ To prevent overfitting, we integrated dropout regularization and an early stopping criterion based on the validation performance. The dataset was partitioned into training, validation, and testing sets (split ratio of 7:1:2). We evaluated the model’s performance using Area Under the Receiver Operating Characteristic curve (AUROC), Area Under the Precision-Recall Curve (AUPRC), and Accuracy.

All computations were performed on a workstation equipped with a single NVIDIA RTX 6000 Ada Generation GPU (48 GB VRAM), a 32-core Intel processor, and 128 GB of RAM. Under this configuration, the full training procedure (100 epochs) was completed in 5.46 hours, which is approximately 1.5 times longer than GCN-CNN (DrugBAN) (3.50 hours). This modest computational requirement, achievable on a single GPU, underscores the efficiency of our GCN-BiLSTM architecture relative to more resource-intensive geometric deep learning models. For the domain adaptation experiments, we evaluated Conditional Adversarial Domain Adaptation (CDAN);^30^ however, our results demonstrated that the baseline GCN-BiLSTM model without domain adaptation delivered superior performance under the current data split.

## 3 Results

### Predictive Performance

We used a curated BindingDB dataset^24^ comprising 49,199 protein-ligand pairs, which encompass 2,623 unique proteins and 14,643 unique ligands, for evaluating the performance of our model. The dataset was partitioned into training, validation, and test sets with an approximate ratio of 7:1:2. Specifically, we used the model trained on the training set to predict the binding probabilities of the unseen pairs in the test set. The performance was rigorously assessed across multiple metrics, including AUROC, AUPRC, accuracy, sensitivity, specificity, and F1-score. The results demonstrate that the proposed GCN-BiLSTM architecture achieves high prediction accuracy, represented by an AUROC of 0.96 and an AUPRC of 0.95. To examine the contribution of the protein BiLSTM representation, we evaluated our model against several alternative architectures by varying the combinations of ligand and protein representations: GCN-CNN (DrugBAN), BiLSTM-BiLSTM, BiLSTM-CNN, and feature-integrated variants (Fig. 2). The proposed GCN-BiLSTM architecture generally outperformed the original CNN-based DrugBAN baseline (AUROC 0.96) and sequence-only variants. These results highlight a distinct hierarchy in representation learning: while GCNs efficiently capture the topological connectivity of small molecules,^16,23^ the protein modality benefits more from the global contextual memory of BiLSTMs. This suggests that modeling long-range sequential dependencies, as a proxy for spatial folding, provides a more robust approximation of the binding context than the restricted local receptive fields of CNNs, especially without explicit 3D structural data.^22^

**Fig. 2.**
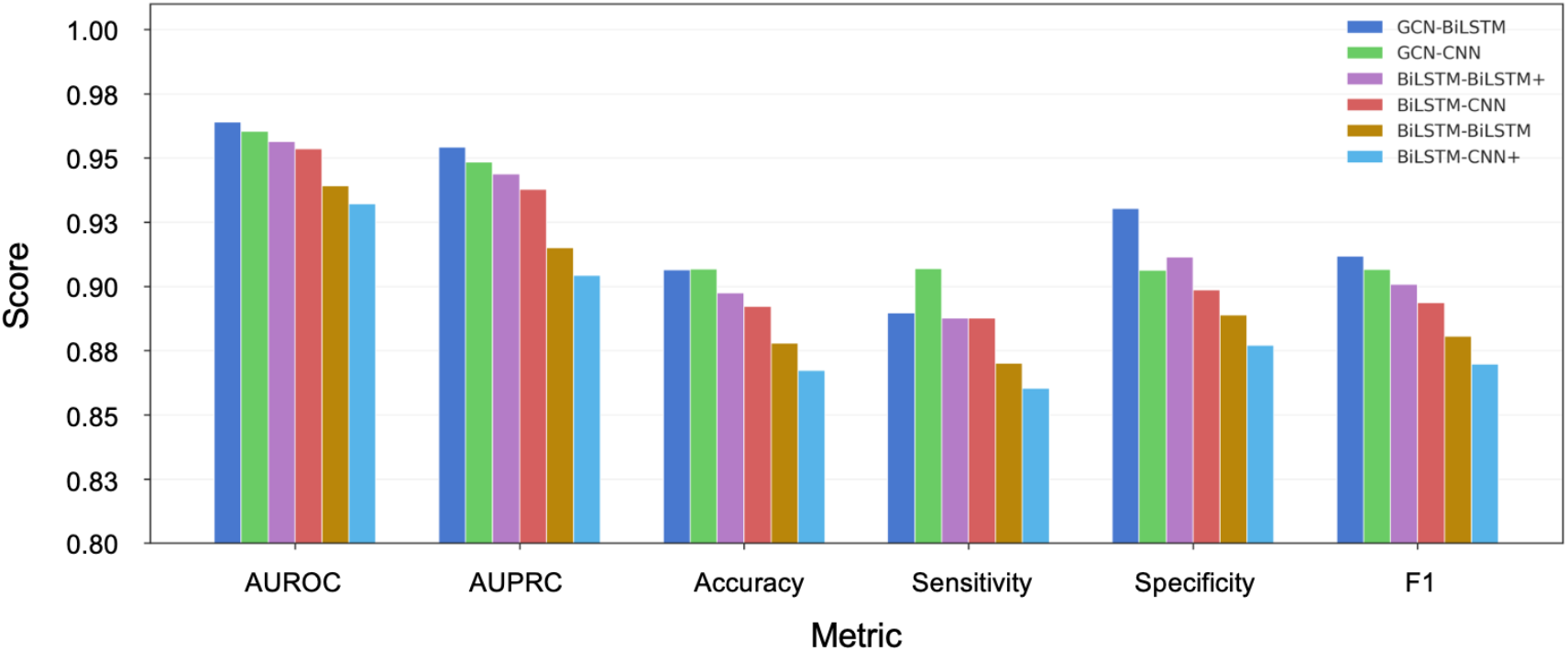
Prediction performance across model architectures. Bar chart comparing six models evaluated on six metrics (AUROC, AUPRC, Accuracy, Sensitivity, Specificity, and F1 score). Models vary in drug and protein encoder types: GCN-CNN (original DrugBAN baseline), GCN-BiLSTM (BiLSTM protein encoder), BiLSTM-CNN and BiLSTM-BiLSTM (fully sequence-based encoders). Feature-augmented variants (denoted with +) additionally incorporate drug physicochemical descriptors (MW, logP, TPSA, hydrogen bond donors/acceptors, ring count, rotatable bonds, and formal charge) and, for BiLSTM-BiLSTM+, per-residue amino acid properties (hydrophobicity, volume, charge, and polarity). All models were trained and evaluated under identical conditions using random data splitting.

### Analysis of Domain Adaptation

Domain shift poses a significant challenge in protein-ligand binding prediction, as models often struggle to generalize across chemically diverse compound spaces. To address this issue, DrugBAN incorporates the conditional adversarial domain adaptation (CDAN),^4,30^ which aligns the feature distributions of different molecule groups. By penalizing domain-specific biases, this approach enables the model to maintain high prediction accuracy for cross-domain protein-ligand binding predictions.

To examine whether the domain adaptations improve our prediction, we evaluated three domain adaptation strategies, CDAN^30^, domain-adversarial neural networks (DANN)^31^, and maximum mean discrepancy (MMD)^32^ to align feature distributions between training and testing clusters. As summarized in Table 1 and Supplementary Fig. S11, these strategies did not consistently improve the proposed GCN-BiLSTM framework (AUROC value of 0.64). Note that CDAN improves the performance of GCN-CNN (DrugBAN) from 0.55 to 0.61 consistent with the previous report.^4^ These results suggest that for the BindingDB cluster-based split, the global protein sequence features extracted by the BiLSTM are inherently robust. Adversarial alignment may impose excessive regularization, potentially hindering the acquisition of fine-grained binding rules.^4,30^ Consequently, we selected the unregularized GCN-BiLSTM as our final architecture.

**Table 1.** Comparison of prediction performance (AUROC values) across various combinations of ligand and protein representations with different domain adaptation strategies: No domain adaptation (Non-DA), the conditional adversarial domain adaptation (CDAN)^30^, the domain-adversarial neural networks (DANN)^31^, and the maximum mean discrepancy (MMD)^32^.

| Encoder Architecture |  | Non-DA<br>(Baseline) | CDAN | DANN | MMD |
| --- | --- | --- | --- | --- | --- |
| Ligand | Protein |  |  |  |  |
| GCN | CNN | 0.55 | 0.61 | 0.59 | 0.55 |
| GCN | BiLSTM | **0.64** | **0.64** | 0.62 | **0.64** |
| BiLSTM | CNN | 0.59 | 0.55 | - | - |
| BiLSTM | BiLSTM | 0.60 | **0.65** | - | - |

### Attention Weight for Interpretation

The advantage of the BAN module lies in its inherent interpretability; it acts as an informational bottleneck, compelling the model to prioritize residues that contribute most significantly to protein-ligand binding.^4,19^ To demonstrate how the model distills robust binding signals from complex data, we visualized the transition from raw attention weights to the final consensus profile (Figs. 3a and 3b, Supplementary Figs. S1–S7). As shown in Fig. 3a, the raw bilinear attention map for target 5HLS (top-scored) reveals dispersed and variable activation patterns across individual ligands. This variability reflects the noise inherent in learning geometric constraints solely from scalar affinity labels. However, our “one-target-multiple-ligands” aggregation strategy effectively filters out these ligand-specific fluctuations. As illustrated in Fig. 3b, this statistical standardization allows the underlying binding signal to emerge clearly from the background noise, causing scattered attention weights to converge into distinct peaks.

**Fig. 3.**
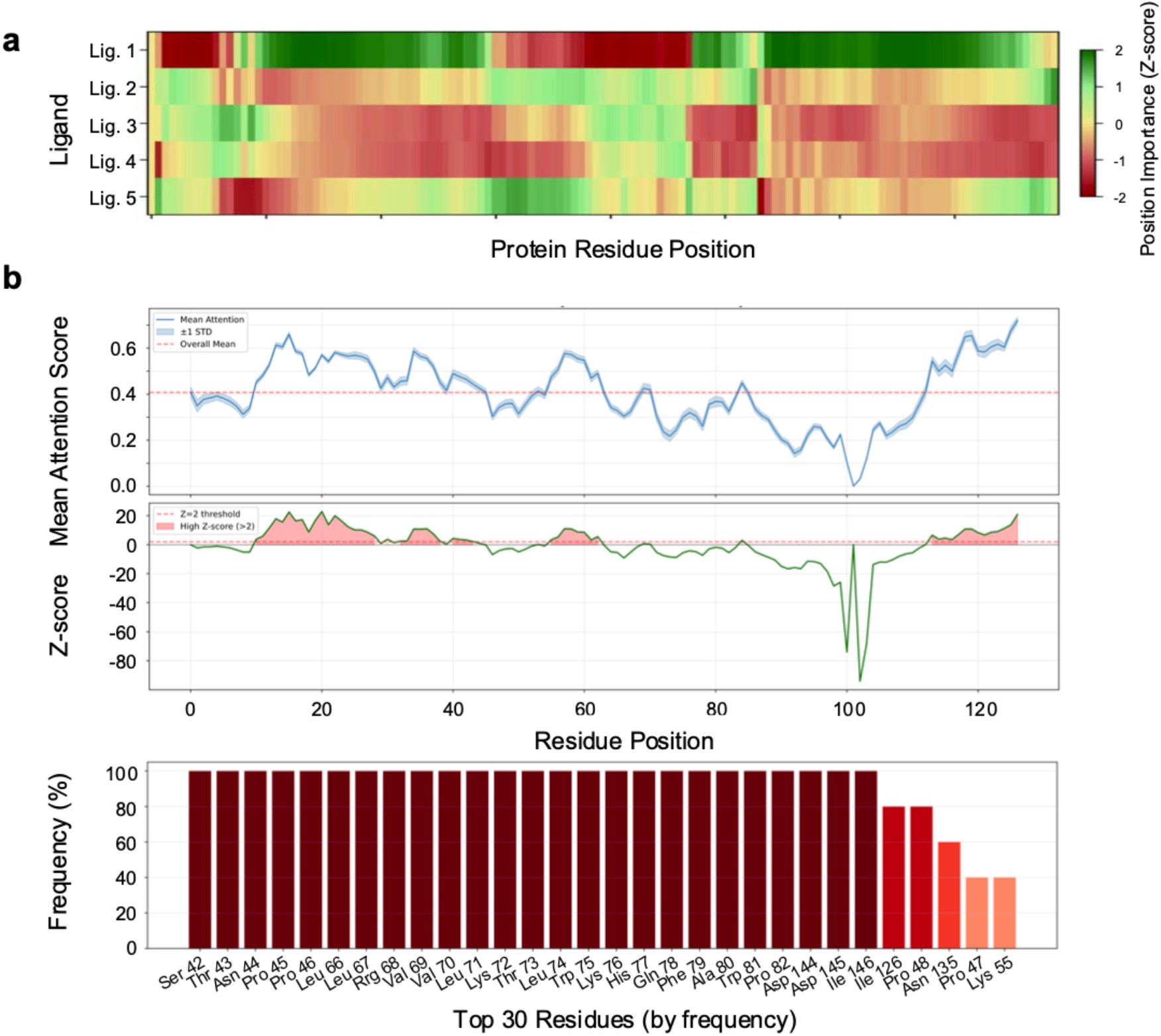
Attention-based binding site profiling. (a) A heatmap representation of raw attention weights across protein residues (x-axis) for five arbitrarily chosen ligands (y-axis), using 5HLS protein (Bromodomain) as an example. We selected five ligands just for illustration purpose. (b) Consensus binding profiles derived from “one-target-multiple-ligands” aggregation: aggregated attention scores (top), standardized Z-scores (middle), and the frequency of residues ranked as top across all ligands (bottom). The ligand-specific noise is effectively filtered to localizes critical binding residues, where residues with high Z-scores (Z > 2) are highlighted in pink.

To evaluate the structural relevance of the attention weights, we systematically compared the predicted binding sites, defined as residues exceeding the 75th percentile of attention scores, against experimentally determined binding pockets from the PDBbind database for eight diverse proteins (Table 2). For the bromodomain 5HLS,^33^ the model identified 27 candidate residues, of which 9 directly overlapped with the ground-truth pocket residues (within 4.0 Å of the ligand), yielding a precision of 33.3%, and a recall of 28.1%. Similarly, for 3NZC,^34^ the model achieved a precision of 33.3% and a recall of 45.5%, demonstrating its ability to recover high-resolution structural insights solely from sequence data. Across the eight target proteins examined, the average precision of 13.50% represents a significant enrichment, approximately 3-5 times higher than random selection (ca. 1–5% for typical pocket-to-sequence ratios). This statistical evidence confirms that the attention mechanism functions as an effective spatial filter, narrowing the search space for downstream structural analysis with greater reliability than qualitative inspections.

**Table 2.** Accuracy of binding site (pocket) localization. The precision of predicted pocket residues overlapping with ground truth was listed for eight diverse proteins. Predicted residues were defined as those with attention weights exceeding the 75th percentile, while ground truth residues were identified based on the PDBbind criteria (residues within 4.0 Å of the ligand). For comparison, the precision of randomly selected residues typically ranges from 1% to –5%.

| PDB ID | Number of pocket residues |  | Precision (%) | Recall (%) | F1-Score (%) |
| --- | --- | --- | --- | --- | --- |
|  | Predicted | Ground truth |  |  |  |
| 3NZC | 30 | 22 | 33.3 | 45.5 | 38.5 |
| 5HLS | 27 | 32 | 33.3 | 28.1 | 30.5 |
| 2QK8 | 22 | 20 | 18.2 | 20.0 | 19.1 |
| 6G3Q | 57 | 19 | 8.8 | 26.3 | 13.2 |
| 1T48 | 39 | 18 | 7.7 | 16.7 | 10.5 |
| 1AQ1 | 44 | 30 | 4.6 | 6.7 | 5.4 |
| 1OXG | 46 | 21 | 2.2 | 4.7 | 3.0 |
| 1QFS | 109 | 20 | 0.0 | 0.0 | 0.0 |
| Average | 46.8 | 22.8 | 13.5 | 18.5 | 15.0 |

### Visualizing Potential Binding Sites

To further validate our model’s interpretability, we examined whether the learned attention weights align with the physical binding site. Visual analysis confirms that the residues assigned high importance scores are consistently localized in the immediate vicinity of the bound ligand (Figs. 4a and 4c). This spatial correlation indicates that the model successfully identifies the critical contact residues necessary for protein-ligand recognition, moving simple affinity prediction toward structural awareness.

**Fig. 4.**
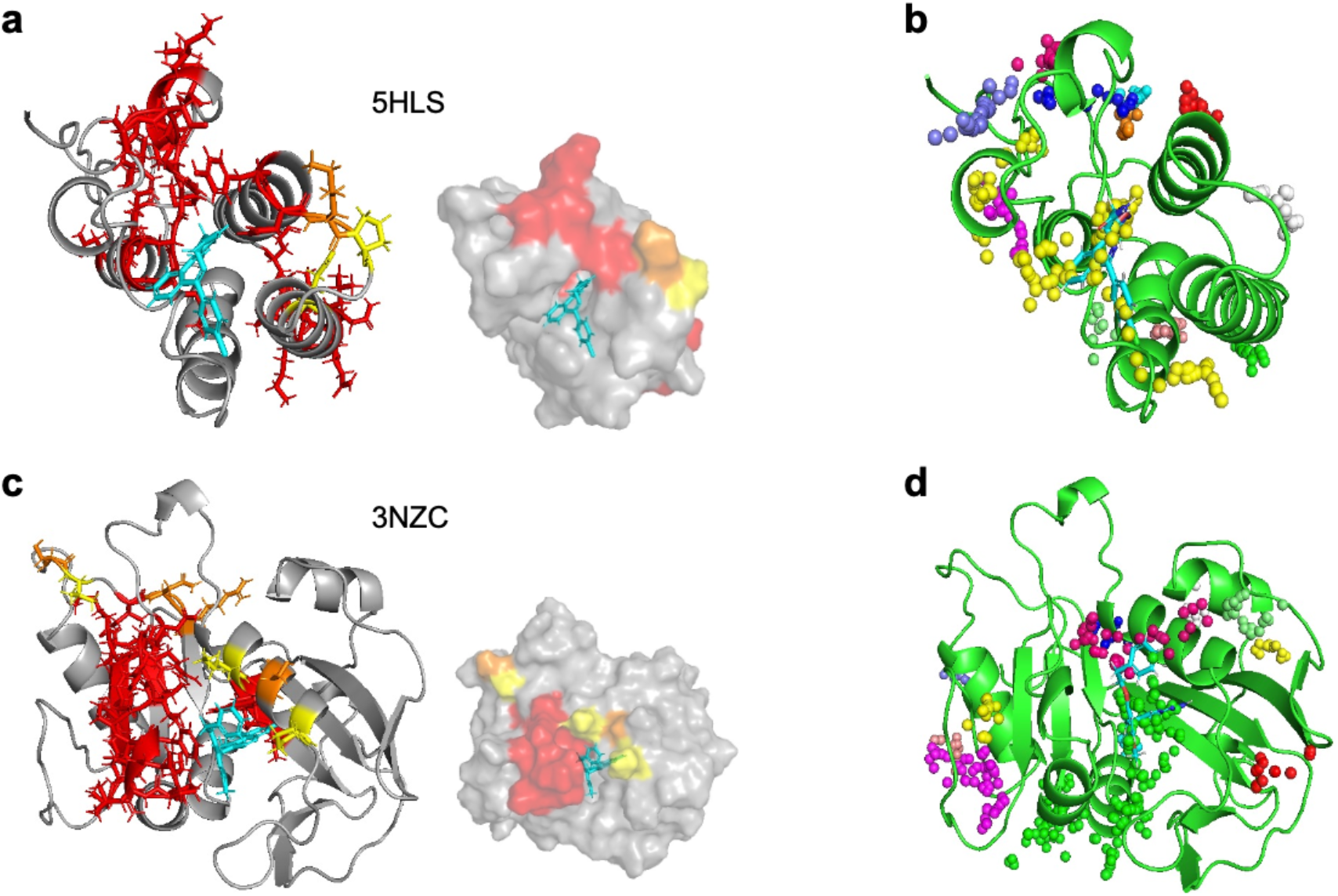
Predicted binding sites. Structures of potential binding pockets for bromodomain 5HLS (top) and dihydrofolate reductase 3NZC (bottom) predicted using the GCN-BiLSTM framework: (a, c) Ribbon and surface representations with bound ligands shown in cyan. The attention frequency is indicated by colors: red (>75%), orange (50–75%), and yellow (25–50%). (b, d) Corresponding binding sites predicted by fpocket (a geometry-based method). Colored spheres represent alpha-spheres clusters filling identified cavities; different colors correspond to distinct candidate pockets.

To evaluate the specificity of these predictions, we compared our results with fpocket,^35^ a widely used geometry-based algorithm that identifies cavities by filling geometric concavities with alpha-sphere clusters. As illustrated in Figs. 4b and 4d, fpocket effectively delineates accessible surface volumes, but it is inherently blind to biological function. Consequently, it often outputs multiple candidate pockets (represented by distinct color clusters in Figs. 4b and 4d) that require manual filtering or docking for verification. In contrast, our GCN-BiLSTM framework acts as a functional filter. While fpocket suggests multiple dispersed cavities, our model’s attention weights tend to converge at the true ligand-binding interface. In other words, the model captures the “chemical semantics” of the binding site, representing residues evolutionarily and chemically primed for interaction, rather than merely identifying geometric voids.

To provide a granular view of the binding interface, we stratified the predicted residues into a three-tiered hierarchy based on their selection frequency across the ligand set: high-confidence (>75%, red), moderate-frequency (50–75%, orange), and lower-frequency (25–50%, yellow). The spatial co-localization of these three tiers within the concave regions suggests a distinct functional division: while the high-confidence (red) residues likely serve as conserved anchor points essential for general binding stability, the moderate and lower frequency (yellow and orange) residues may drive ligand-specific selectivity, interacting only with distinct molecular substructures. This hierarchical convergence is not unique to our primary case studies but is consistently observed across the broader validation set (e.g., 6G3Q, 1T48, and 1QFS; Supplementary Figs. S8–S10), demonstrating that the model’s ability to prioritize core binding anchors is a robust and generalized feature of its learning architecture.

### Comparison with CNN-based Baseline (DrugBAN)

To rigorously benchmark our model against the CNN-based baseline (DrugBAN), we evaluated the predicted binding sites for three distinct receptors (4N6H,^36^ 5W8L,^37^ and 6QL2^38^) featured in the original study.^4^ The comparative performance highlights that the optimal encoder architecture is intrinsically linked to the specific topological characteristics of the protein target (Table 3).

**Table 3.** Comparison of binding site prediction accuracy: BiLSTM vs. CNN baseline. The number of correctly identified binding site (pocket) residues (Recall) is shown for three representative receptors (delta-type opioid receptor (4N6H), L-lactate dehydrogenase (5W8L), and carbonic anhydrase (6QL2)). Results are categorized by attention frequency: >75%, 50–75%, and 25–50%. Ground truth residues were defined as those within 4.0 Å of the ligand in the corresponding PDB structure.

| Receptor | Ligand | Number of pocket residues |  | Recall (%) |
| --- | --- | --- | --- | --- |
|  |  | Ground truth | Predicted<br>>75%/50–75%/25–50%<br>(Protein encoder) |  |
| 4N6H | EJ4 | 11 | 2/0/0 (CNN) | 18.2 |
|  |  |  | 2/0/0 (BiLSTM) | 18.2 |
| 5W8L | 9YA | 16 | 1/0/0 (CNN) | 6.2 |
|  |  |  | 6/0/1 (BiLSTM) | 43.8 |
| 6QL2 | EZL | 10 | 5/0/0 (CNN) | 50.0 |
|  |  |  | 0/1/0 (BiLSTM) | 10.0 |

For the target 5W8L (9YA ligand bound to human L-lactate dehydrogenase A), our BiLSTM-based model demonstrated a substantial advantage, achieving a recall of 43.8% compared to a mere 6.2% for the CNN baseline. Visual analysis (Figs. 5a and 5b) confirms that the BiLSTM-derived attention weights successfully form a spatially coherent cluster around the ligand (9YA), whereas the CNN model identifies only a single relevant residue and fails to define a pocket. Interestingly, for the target 4N6H (human delta-type opioid receptor with EJ4 ligand), both the BiLSTM and CNN encoders exhibited identical baseline performance, each achieving a recall of 18.2% by correctly identifying two high-confidence residues. Conversely, for target 6QL2 (ethoxzolamide (EZL) complexed with human carbonic anhydrase 2), the CNN baseline outperformed the BiLSTM model (50.0% vs. 10.0% recall), successfully capturing specific local motifs proximal to the ligand (EZL) (Figs. 5a and 5b).

**Fig. 5.**
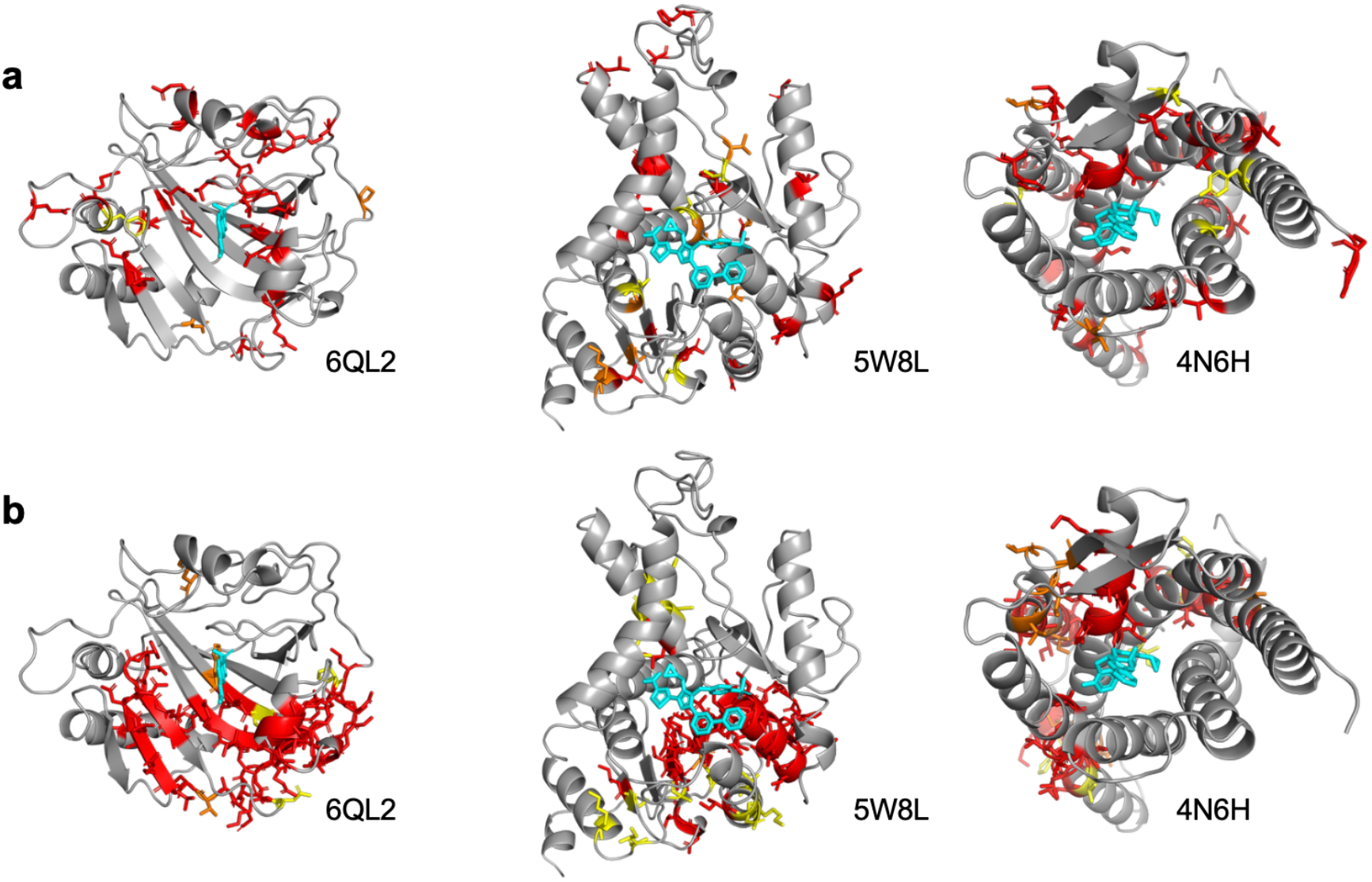
Comparison of predicted binding sites: BiLSTM vs. CNN baseline. Predicted potential binding sites for three distinct receptors (4N6H, 5W8L, and 6QL2) originally reported^4^, comparing (a) GCN-CNN (DrugBAN) and (b) GCN-BiLSTM frameworks. The attention frequency is indicated by colors: red (>75%), orange (50–75%), and yellow (25–50%). The bound ligands are shown in cyan.

This divergent performance can be directly attributed to the distinct geometric and biochemical characteristics of the binding pockets. For 5W8L (L-lactate dehydrogenase A), the binding site is relatively large and dynamic, interacting with both a coenzyme (NADH) and a ligand across a broad protein interface. This suggests that the BiLSTM’s ability to capture global sequence context and long-range dependencies is essential for defining such multicomponent pockets. In contrast, 6QL2 (carbonic anhydrase 2) features a localized, rigid, and narrow conical pocket centered around a zinc coordination site. For such compact sites defined by distinct, contiguous local motifs, the localized receptive fields of CNN convolutional filters may more efficiently identify “lock-and-key” signatures than recurrent architectures.

Analysis of the sequence-level attention profiles further corroborates this hypothesis. The CNN encoder produces high-frequency, discontinuous spikes scattered across the sequence (Supplementary Fig. S12-S19). This fragmentation indicates that the CNN extracts isolated motifs without integrating the broader sequential context, resulting in a dispersed spatial distribution that often fails to delineate a coherent binding pocket in 3D space (Supplementary Fig. S20-S22). In contrast, the BiLSTM-derived profiles demonstrate smoother, continuous peaks that translate into highly localized clusters at the actual binding interface. Ultimately, these findings indicate that while the BiLSTM offers broader robustness for complex and dynamic interfaces, the efficacy of the encoder architecture is highly dependent on the topological complexity of the target pocket.

## 4 Discussion

Quantitative evaluation across eight test proteins yielded an average precision of 13.50% and a recall of 18.50% (Table 2). While these sequence-level metrics may appear modest compared to 3D-supervised methods, they offer substantial utility for structural screening. Given that a typical binding pocket occupies only 5-10% of 500-1000 residue sequences, our model effectively narrows the search space from the entire protein surface to high-priority regions. Rather than performing exhaustive blind docking, one can focus intensive physics-based sampling on these predicted hotspots by “Attention GPS”. The incorporation of our GCN-BiLSTM attention map into the existing structural pocket analysis facilitates the functional characterization of specific sites that are otherwise difficult to identify. For instance, fpocket effectively identifies accessible volumes but lacks the biological context to distinguish active sites from inert depressions.^35^ By applying the BiLSTM attention as a functional filter, we successfully eliminated numerous false-positive geometric voids, narrowing the search space to the crystallized orthosteric interface (visualized in magenta, Fig. 6a and Supplementary Fig. S23). Intriguingly, the regions captured by our model in 5HLS demonstrate a striking consensus with known allosteric sites (Fig. 6b).^39^ This alignment is particularly significant given that the structural flexibility of bromodomains, epigenetic readers recognizing acetylated lysines, is fundamental to their functional control.^40^ This observation suggests our model’s effectiveness even in identifying the allosteric regulatory elements and conformational plasticity.

**Fig. 6.**
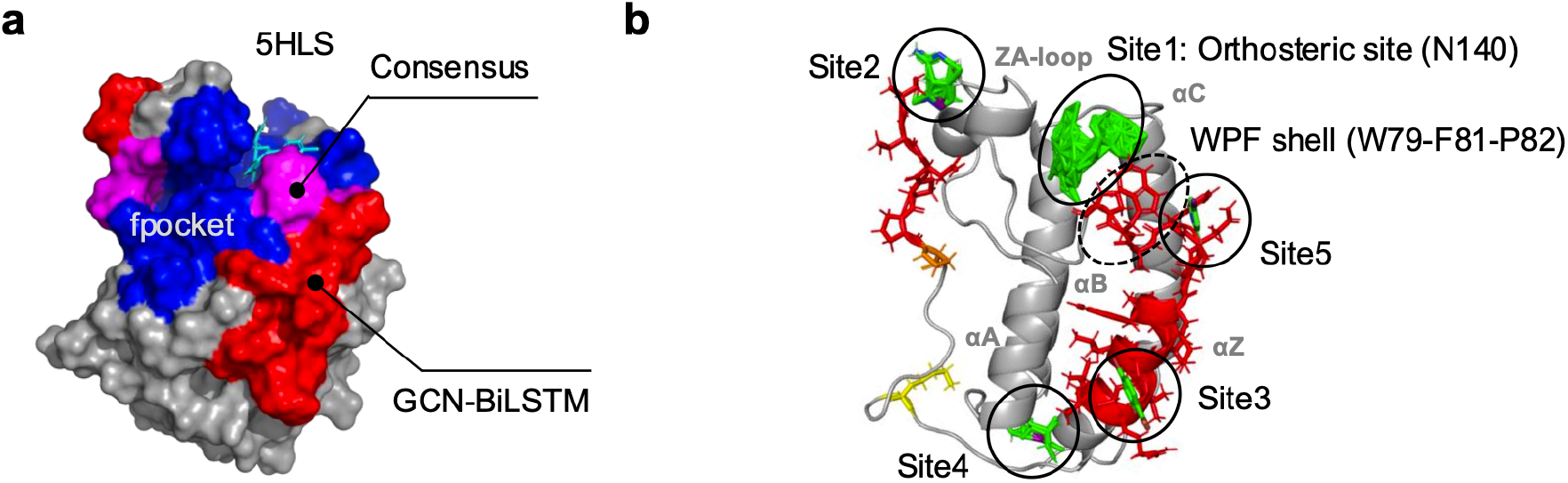
Comparison of geometric (fpocket) with semantic (GCN-BiLSTM) predictions. (a) Surface representations of binding sites predicted for 5HLS: the regions unique to BiLSTM and fpocket are shown in red and blue, respectively. Magenta indicates the intersection of BiLSTM attention and fpocket cavities. (b) Attention frequency mapped onto the 5HLS structure (red: >75%, orange: 50–75%, and yellow: 25–50%). Ligands bound to known orthosteric (Site1) and allosteric sites (Site2-5) (taken from Ref. [39]) are shown for comparison (green). The bound ligand of 5HLS is shown in cyan.

Our analysis also identified critical boundary conditions, most notably in the case of 1QFS, a prolyl oligopeptidase with a deeply buried active site,^41^ which yielded 0.00% precision (Table 2). Structural analysis (Supplementary Fig. S9) reveals that the binding site of 1QFS resides within a central cavity formed by its β-propeller domain, which largely exceeds our sequence encoder’s truncation point (1,200 residues). Rather than a simple failure, this constructive outlier serves as a crucial control: it confirms that the model does not randomly hallucinate binding sites but is strictly constrained by the provided input information. This underscores the necessity of aligning sequence-length capacities with the structural complexity of large, multi-domain proteins.

While AlphaFold3 has set a new benchmark for atomic-level accuracy in biomolecular complex prediction, its generative diffusion process is computationally prohibitive for exhaustive giga-scale chemical library screening.^10^ In this context, our sequence-based framework is designed not to replace high-precision structural modeling, but to serve as a high-throughput “funnel” (pre-screening filter). This hierarchical strategy aligns with the machine learning-boosted docking paradigm,^5,14^ where rapid machine learning models effectively reduce billions of candidates to a manageable subset for rigorous verification. By navigating vast chemical spaces without the overhead of 3D coordinate generation, our GCN-BiLSTM model complements AlphaFold3, prioritizing high-probability leads that can subsequently be refined by computationally intensive structural tools.

## 5 Conclusion

In this study, we developed a weakly supervised deep learning framework that bridges the gap between high-throughput sequence screening and structural interpretability. By integrating GCN and BiLSTM, our architecture overcomes the receptive field limitations of traditional CNNs, effectively capturing the long-range dependencies essential for protein modeling. The resulting model achieves state-of-the-art performance on the BindingDB dataset (AUROC 0.96) and functions as an interpretable “Attention GPS”. Despite the absence of 3D-coordinate supervision, the BAN successfully localizes key binding residues that align with experimental crystal structures.

Our synergistic analysis demonstrates that overlaying sequence-based attention with geometry-based cavity detection (fpocket) identifies a robust consensus zone. This integrated approach effectively filters out inert geometric voids and highlights a dynamic binding landscape, including potential allosteric sites. While challenges remain for ultra-large multi-domain proteins like 1QFS, the framework’s intrinsic generalizability offers a scalable solution for early-stage drug discovery. Ultimately, by transforming computationally expensive blind docking into targeted local exploration, our framework proves that AI-driven chemical intuition and physics-based constraints are highly complementary. This integration provides a high-throughput paradigm that balances speed, accuracy, and mechanistic interpretability for modern structure-based drug design.

## Contributions

C.-Y.C. conceived the study, performed the experiments, and wrote the original draft. F.-Y.L., Y.-A.C., and S.R. provided conceptual guidance and professional expertise throughout the research. All authors contributed to the refinement and revision of the manuscript. S.R., Y.-A.C., and F.-Y.L. supervised the research.

## Acknowledgements

This work was supported by the “Doctoral Study Abroad Scholarship” from National Chung Hsing University. The authors gratefully acknowledge the National Institutes of Biomedical Innovation, Health and Nutrition (NIBN) for providing the computational resources and research facilities essential for the completion of this work.

## Notes

### Competing Interest Statement

The authors have declared no competing interest.

