## Supplementary Information for "BiLSTM-Powered Bilinear Attention for Protein–Ligand Prediction"

**Comprehensive analysis of attention distributions and structural mappings**

To ensure the robustness of our interpretability findings, we extended the visualization analysis to the remaining six proteins in our validation set (in addition to the 5HLS and 3NZC cases presented in the main text). Supplementary Figs. S1–S7 illustrate the raw bilinear attention weight distributions across the protein sequences. These heatmaps reveal that the model does not simply attend to the entire sequence uniformly; instead, it learns sparse, distinct regions of high importance, reflecting specific biochemical motifs rather than background noise.^1^

**Structural validation and error analysis**

We further projected these high-attention residues onto their respective 3D crystallographic structures to assess spatial coherence (Supplementary Figs. S8–S10). Crucially, the structural mappings for targets such as 2QK8 and 6G3Q (Supplementary Fig. S8) corroborate the “functional hierarchy” hypothesis proposed in the main text. We observed a recurrent pattern where high-frequency “anchor” residues (red) cluster centrally near the ligand, flanked by moderate-frequency “specificity” residues (yellow/orange).

However, Supplementary Fig. S9 highlights a critical boundary condition of our current framework. Specifically, for the target 1QFS, the model failed to identify the binding pocket (Precision 0.00%). Structural analysis reveals that 1QFS is a large multi-domain protein complex where the binding site is located deep within the central core or at a sequence position exceeding our model's truncation threshold (1,200 residues). This negative result serves as an important control, confirming that the model's predictions are constrained by the input information content (sequence length coverage) and do not rely on hallucinated structural priors.^1^

**Impact of domain adaptation strategies on predictive accuracy**

We further investigated the utility of explicit domain alignment by integrating Conditional Adversarial Domain Adaptation (CDAN) into our framework.^2^ As illustrated in Supplementary Fig. S11, this strategy led to a substantial decline in predictive performance, with AUROC scores dropping by over 30% across all tested encoder configurations compared to the baseline. This finding suggests that in the context of the BindingDB cluster-based split, adversarial alignment imposes excessive regularization, hindering the model's ability to learn the fine-grained binding rules required for accurate affinity prediction.^1,2^

**Comparative analysis of attention topologies: CNN-based baseline**

To provide a comprehensive benchmark for our interpretability findings, we extended the visualization analysis to the baseline DrugBAN model (utilizing the CNN encoder) across the validation set, as presented in Supplementary Figs. S12–S22. These visualizations facilitate a direct topological comparison with the BiLSTM-derived results shown in Supplementary Figs. S1–S10.

Consistent with the representative cases discussed in the main text (Fig. 5), the CNN baseline exhibits a systematic inability to model sequential continuity. As exemplified by the detailed profiles for 5HLS (Supplementary Fig. S12) and 3NZC (Supplementary Fig. S18), the sequence-level attention distributions are dominated by high-frequency, discontinuous spikes. This contrasts sharply with the smooth, wave-like attention motifs generated by our GCN-BiLSTM framework (Supplementary Figs. S1–S7), which reflect the continuous nature of protein domains. When projected into 3D space, this signal fragmentation manifests as spatially dispersed residue clusters, as seen in the structural comparisons of 6QL2, 5W8L, and 4N6H (Fig. 5a).

When projected into 3D space, this signal fragmentation manifests as spatially dispersed residue clusters. Unlike the BiLSTM model, which identifies consolidated "anchor" regions that tightly envelop the ligand (cyan), the CNN model consistently highlights scattered surface residues that fail to define a coherent binding interface. This pattern is pervasive across the broader dataset (Supplementary Figs. S12–S22), reinforcing the conclusion that the global contextual modeling provided by BiLSTM is essential for overcoming the limitations of local receptive fields in weakly supervised binding site localization.

**Comparison of geometric and semantic pocket predictions: “functional filtering”**

The integration of our GCN-BiLSTM attention map with conventional structural pocket analysis enhances the functional characterization of sites that evade detection by geometry-based methods alone. While fpocket^3^ excels at identifying accessible volumes, it often fails to differentiate biologically active sites from inert surface depressions. By utilizing our attention map as a functional filter, we effectively pruned false-positive geometric voids, narrowing the search space to the crystallized orthosteric interface. This synergistic effect is visually confirmed by the spatial overlap between semantic attention and geometric cavities (Supplementary Fig. S23), where the model successfully filters out non-functional structural voids to highlight the true binding pocket.

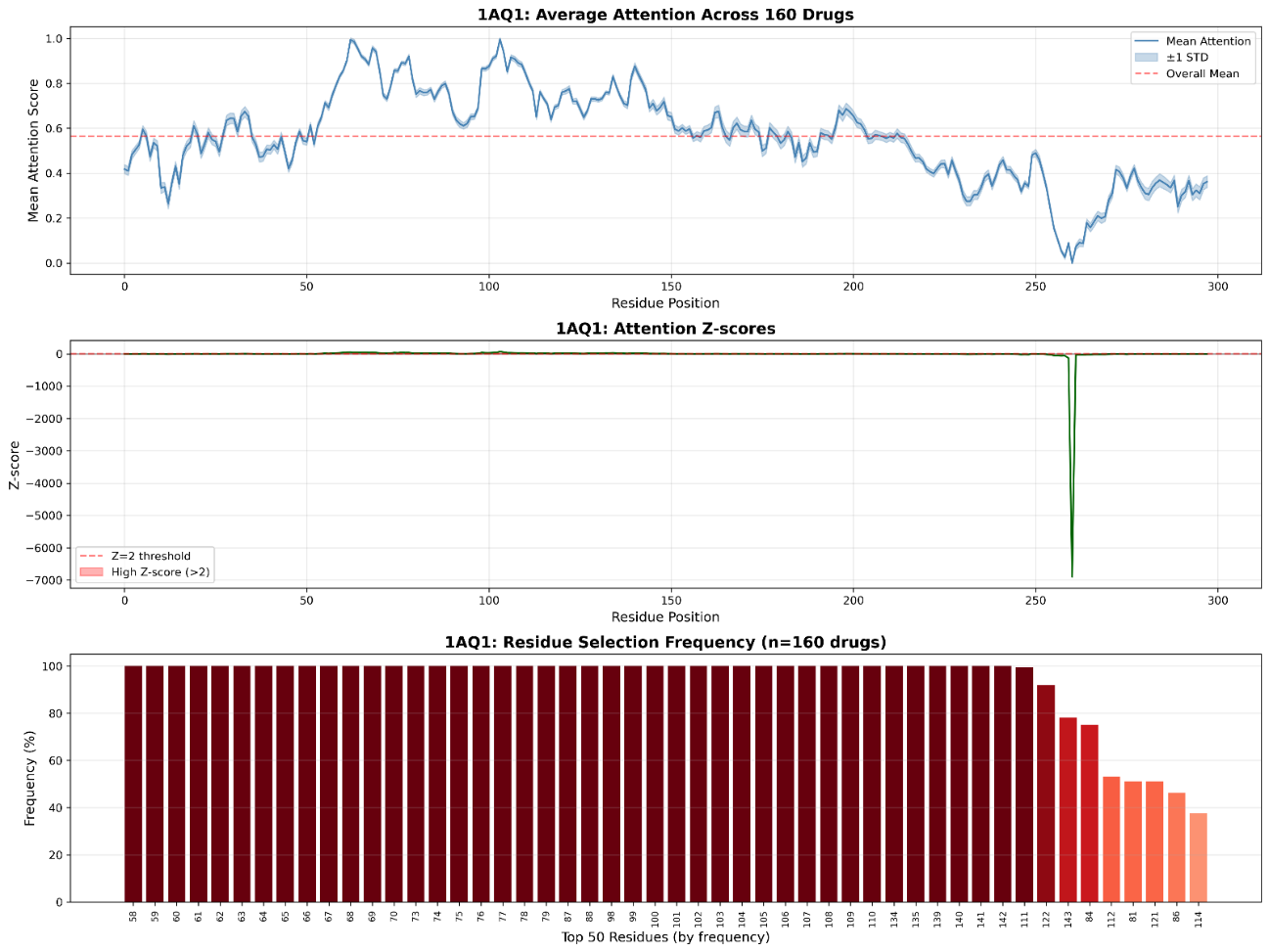

Supplementary Fig. S1 | Learned attention profile for 1AQ1. The visualization displays the mean attention score (top) and standardized Z-scores (middle) across the amino acid sequence, alongside residue selection frequency (bottom). Significant peaks and continuous regions of high attention correspond to residues implicitly identified by the model as critical for interaction.

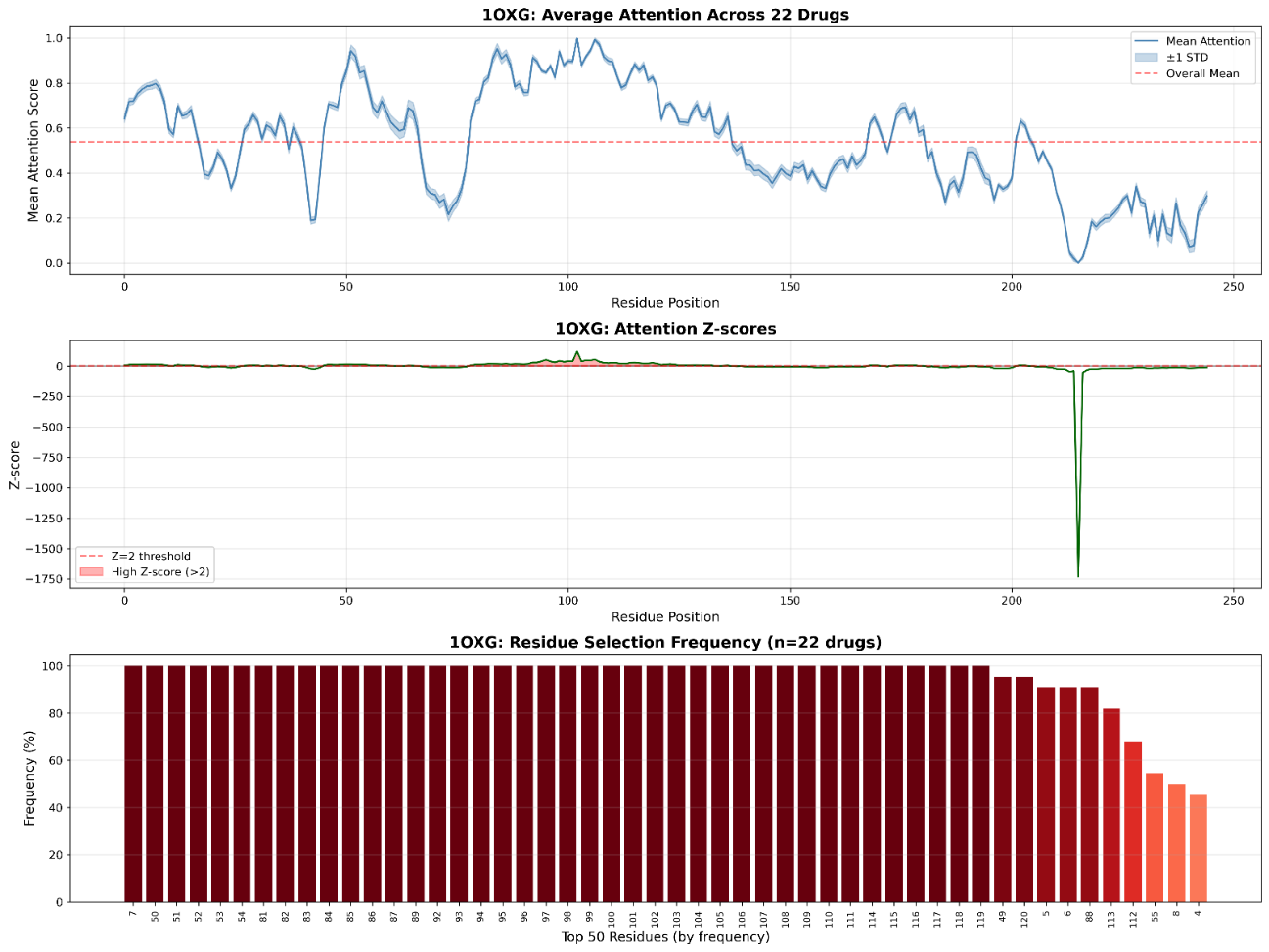

Supplementary Fig. S2 | Learned attention profile for 1OXG. Visualization of the attention statistics for protein 1OXG, showing mean attention (top), Z-scores (middle), and selection frequency (bottom). The sparse distribution of high-intensity regions suggests the model is focusing on specific local motifs rather than global sequence features.

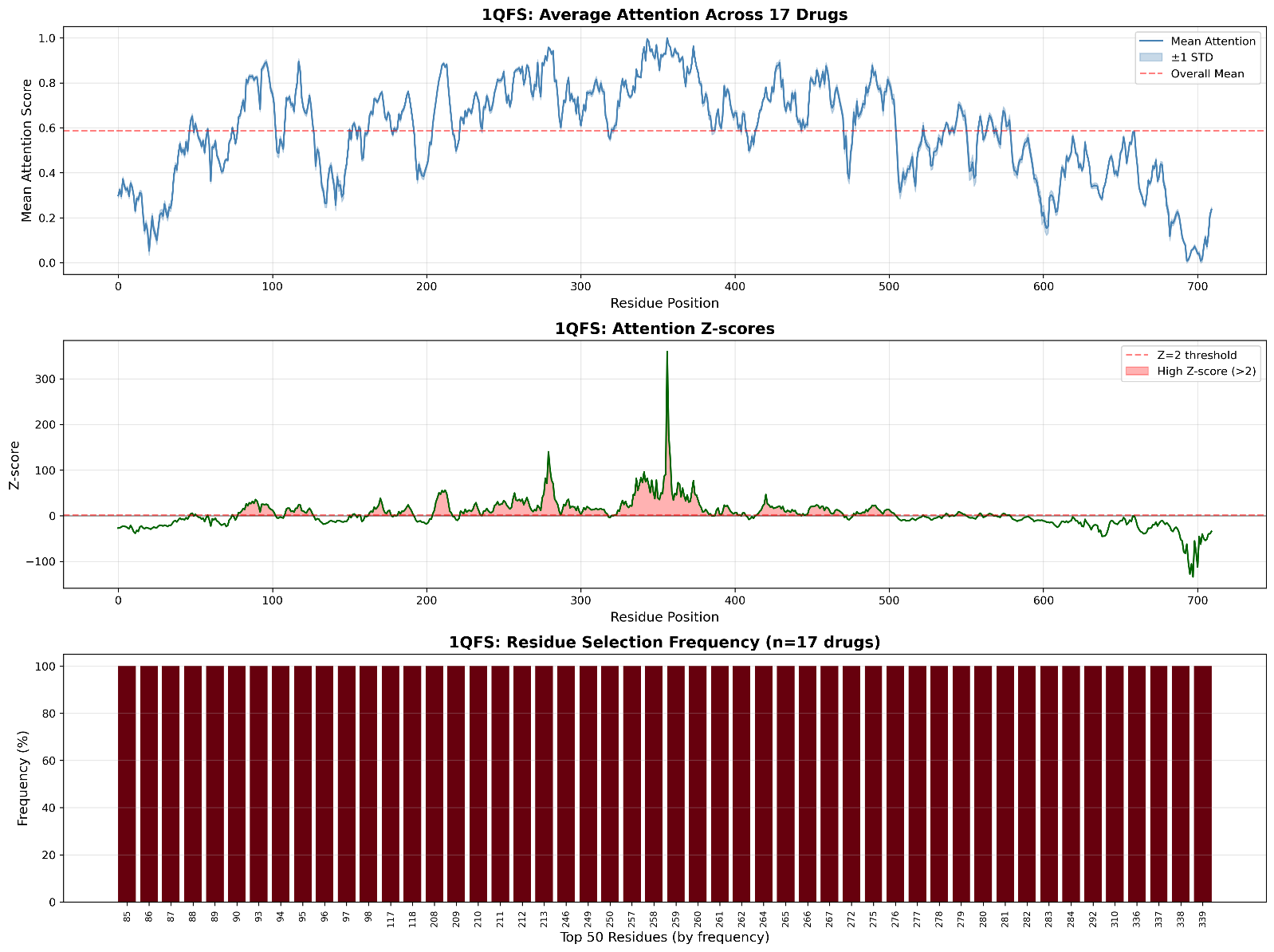

Supplementary Fig. S3 | Learned attention profile for 1QFS. Attention profile for the protein 1QFS displaying mean scores (top), Z-scores (middle), and residue frequency (bottom). Note the complex distribution of peaks; this protein represents a challenging case due to its large sequence length and multi-domain structure, leading to the “constructive failure” analyzed in Supplementary Fig. S9.

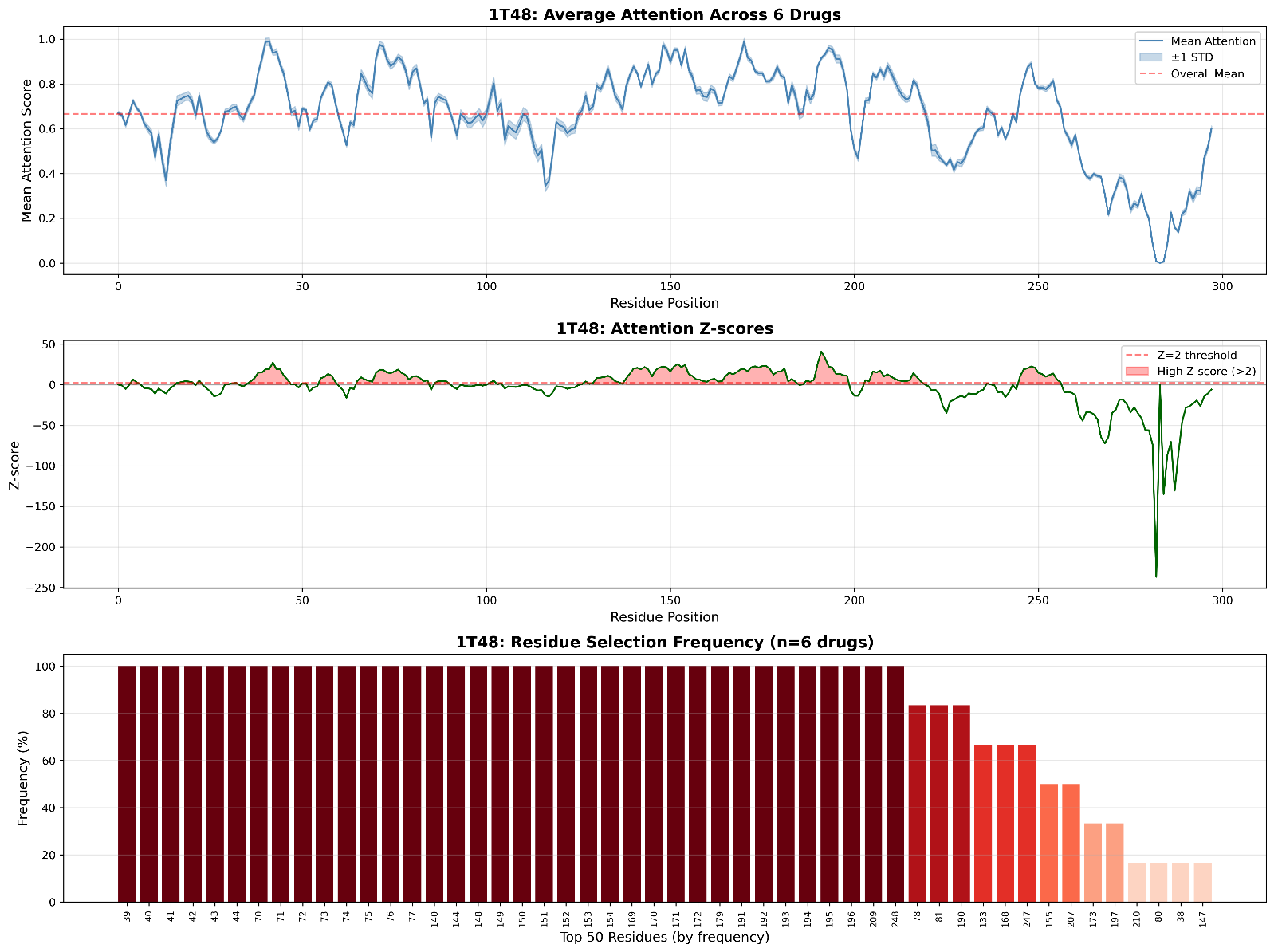

Supplementary Fig. S4 | Learned attention profile for 1T48. Attention profile for 1T48 showing mean attention (top), Z-scores (middle), and frequency (bottom). The model identifies distinct regions of interest along the sequence which act as candidates for the ligand-binding interface.

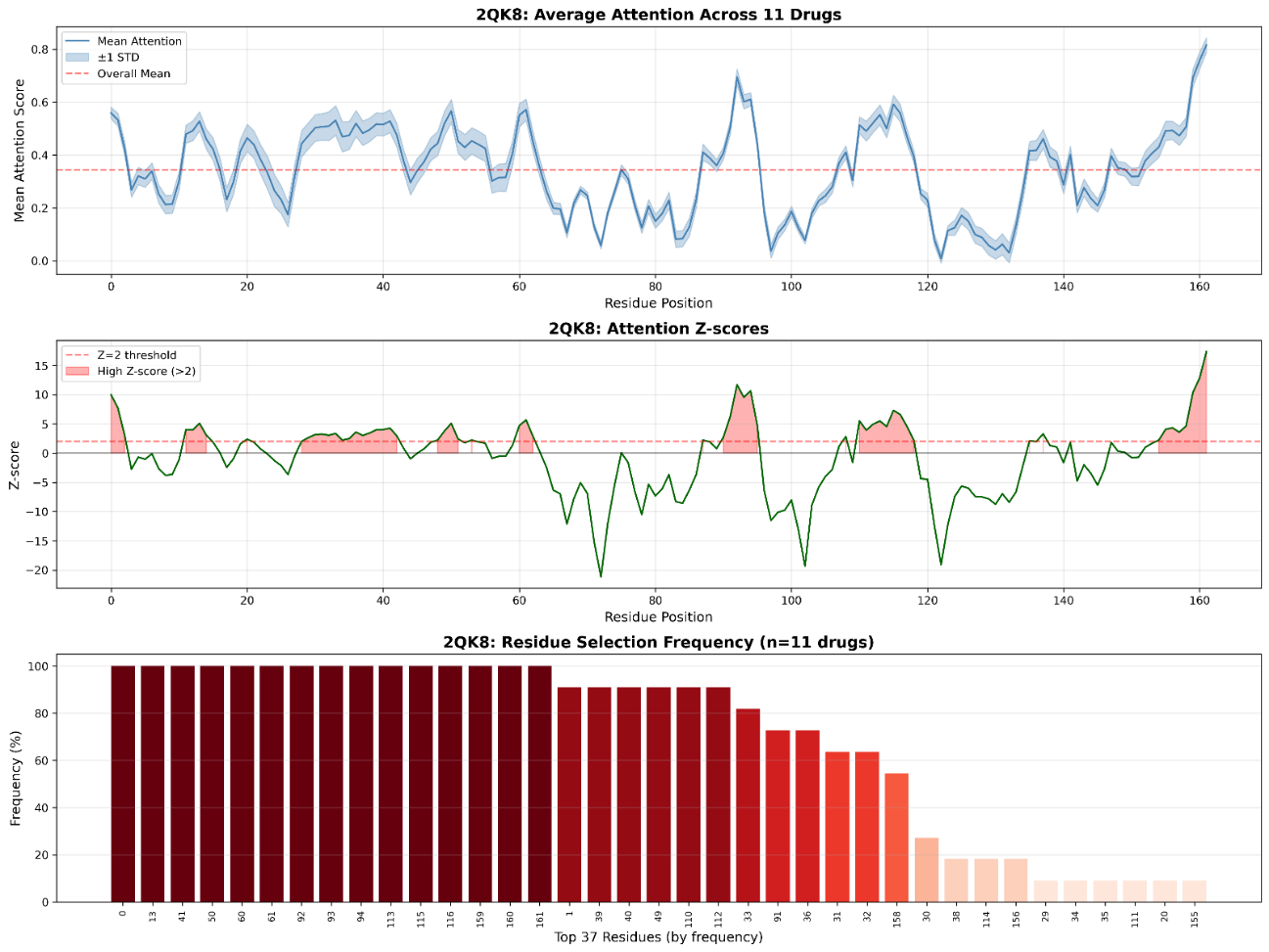

Supplementary Fig. S5 | Learned attention profile for 2QK8. Visualization of attention statistics for 2QK8. The high-frequency residues in the bottom panel (dark red) correspond to the model's predicted “anchor points” for ligand binding, validated structurally in Supplementary Fig. S8.

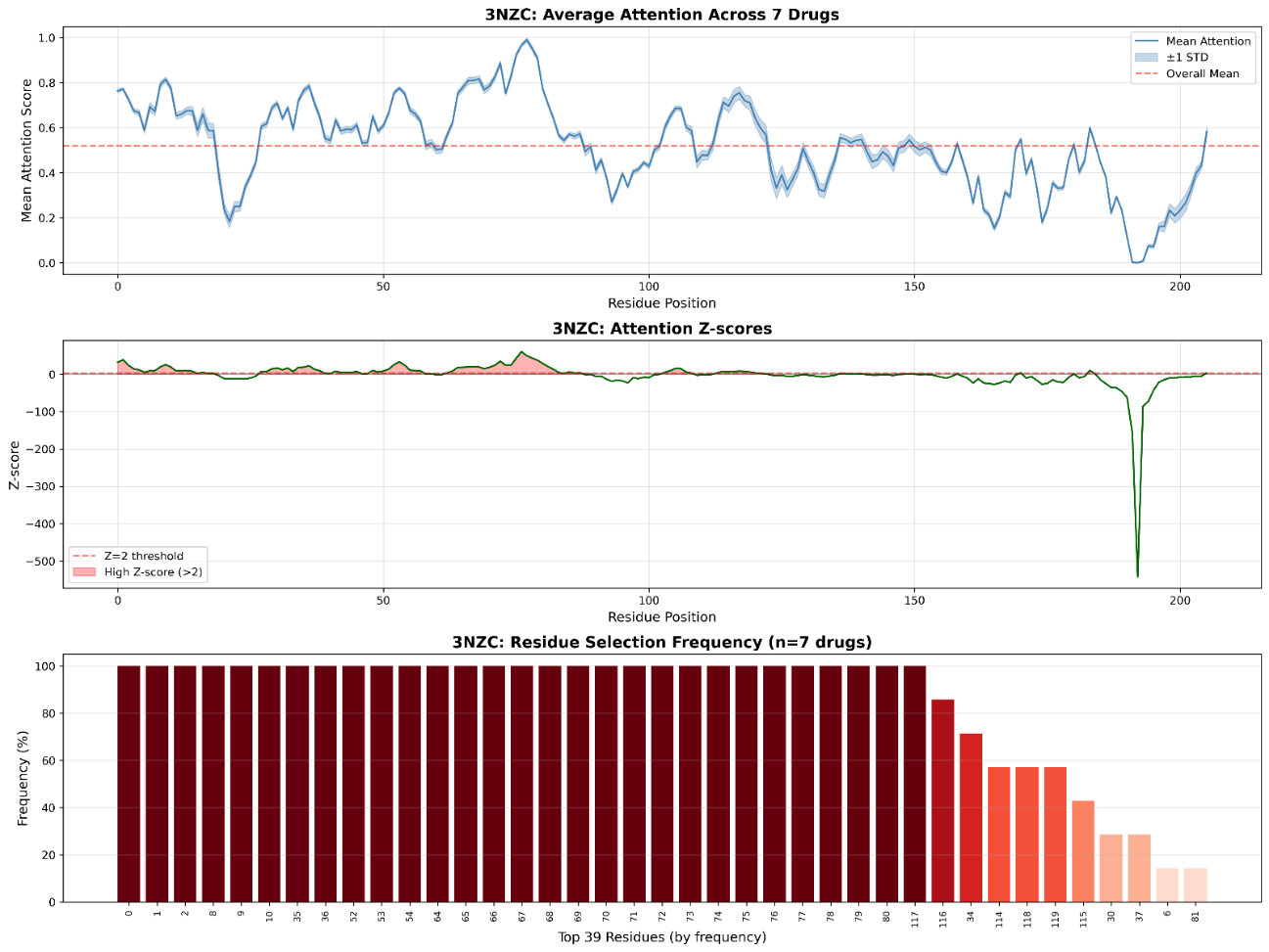

Supplementary Fig. S6 | Learned attention profile for 3NZC. Detailed attention profile for 3NZC (dihydrofolate reductase). This sequence-level visualization corresponds to the successful structural prediction shown in the main text (Fig. 4c), displaying sharp, well-defined attention peaks in the Z-score plot (middle) and high consistency in residue selection frequency (bottom). The convergence of attention on specific residues provides the mechanistic basis for the accurate localization of the orthosteric binding pocket observed in the 3D model.

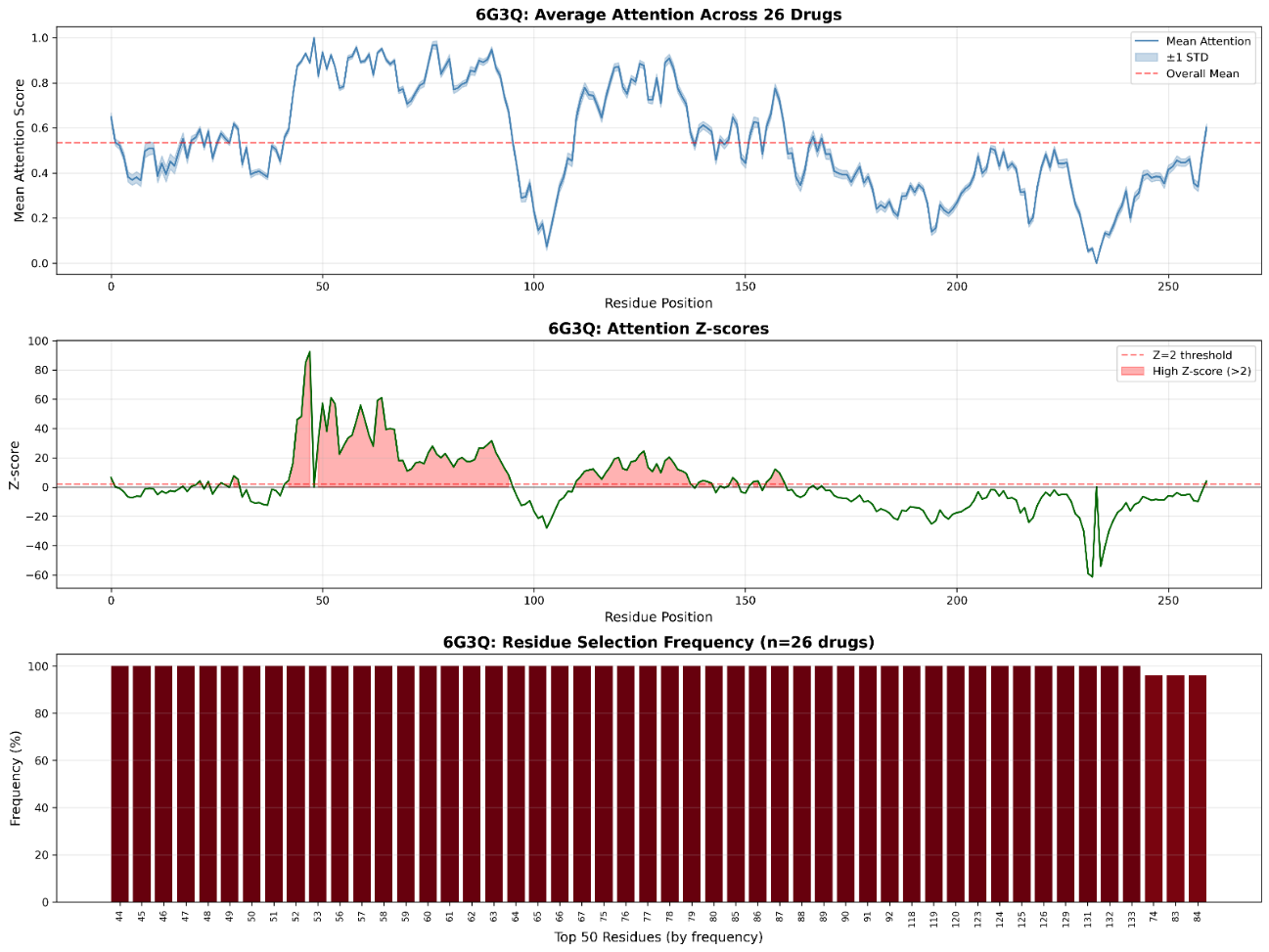

Supplementary Fig. S7 | Learned attention profile for 6G3Q. Attention weight distribution for 6G3Q. The model highlights specific residue clusters in the frequency plot (bottom) that are evaluated for spatial proximity to the ligand in Supplementary Fig. S8.

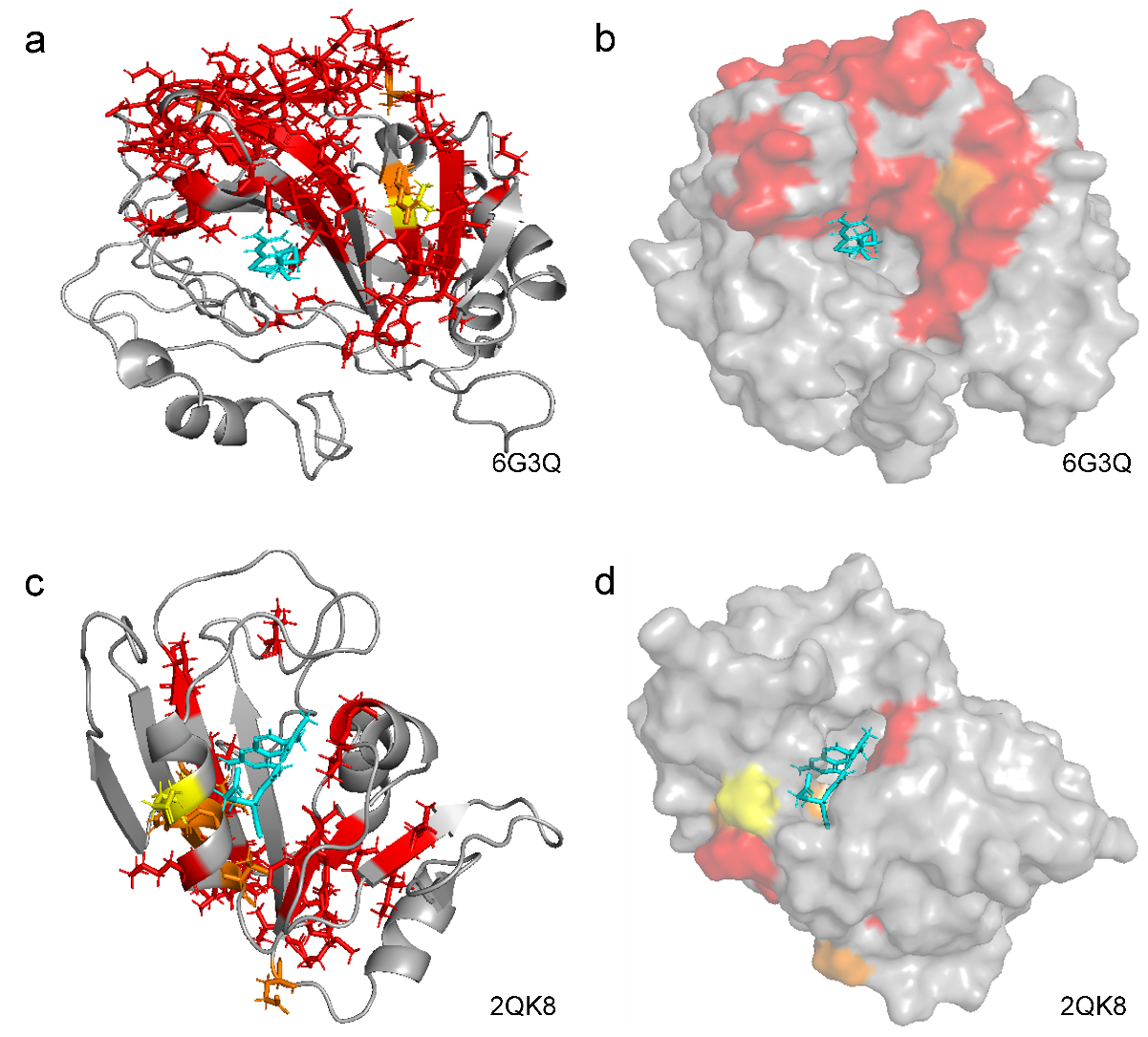

Supplementary Fig. S8 | Structural validation of binding site localization for 6G3Q and 2QK8. 3D molecular visualizations of 6G3Q (a, b) and 2QK8 (c, d). Panels (a) and (c) display ribbon representations, while panels (b) and (d) show surface representations. Residues are colored by attention frequency: red (>75%), orange (50–75%), and yellow (25–50%). Consistent with the main text, the red high-confidence residues cluster centrally around the ligand (cyan), flanked by orange/yellow specificity determinants. This demonstrates the model's robust ability to localize binding regions across diverse protein folds.

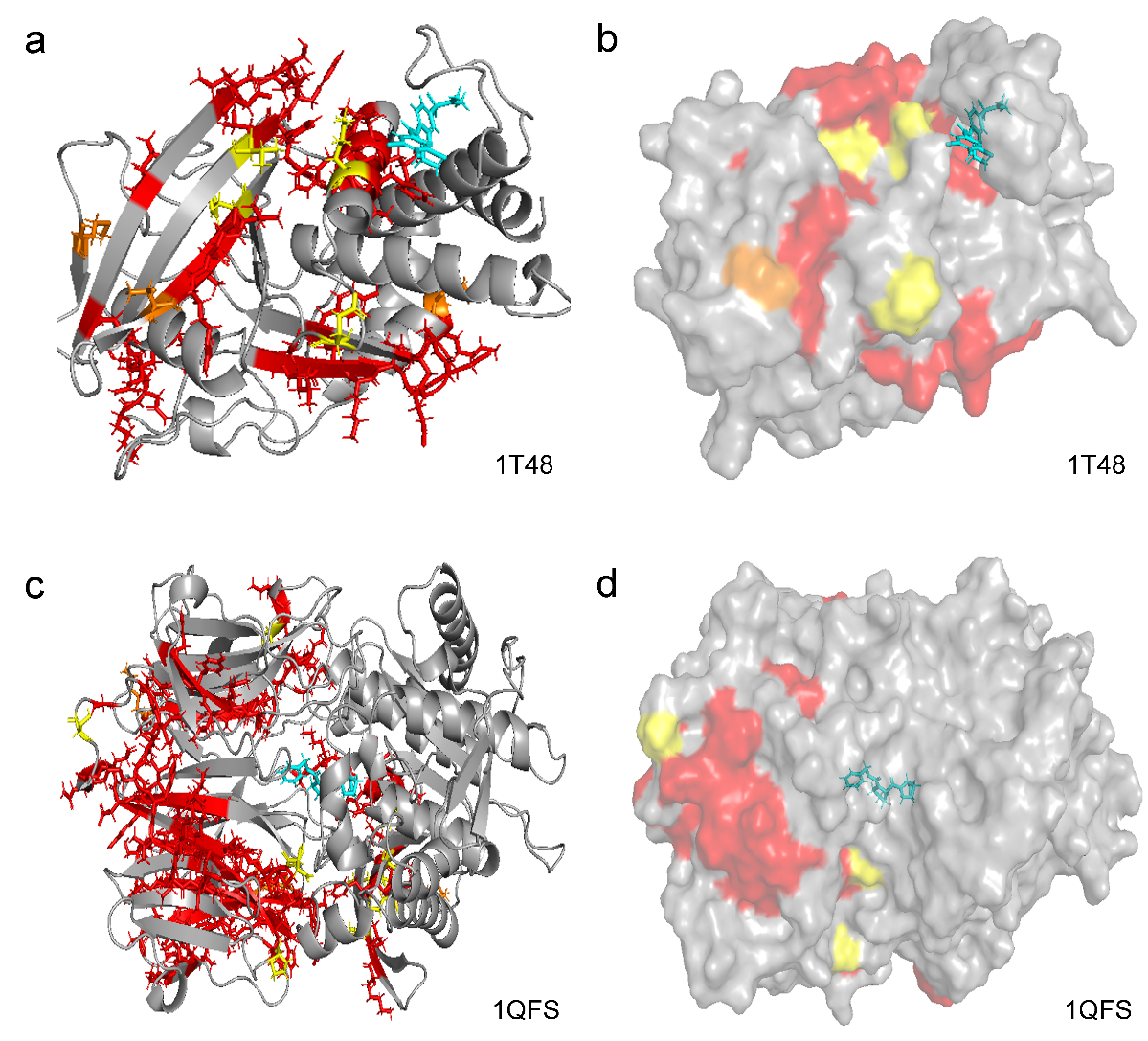

Supplementary Fig. S9 | Structural validation of binding site localization for 1T48 and 1QFS. 3D molecular visualizations of 1T48 (a, b) and 1QFS (c, d). Panels (a) and (c) display ribbon representations, while panels (b) and (d) show surface representations. Residues are colored by attention frequency: red (>75%), orange (50–75%), and yellow (25–50%). For 1T48 (a, b), the high-confidence residues (red) exhibit partial spatial convergence, localizing in general proximity to the ligand (cyan) alongside some distal secondary structures. Conversely, 1QFS (c, d) serves as a critical negative control. The predicted high-frequency regions (red) are densely clustered on a distal domain, completely failing to localize around the true binding interface (cyan). This negative structural result visually confirms the boundary conditions of the current framework when processing large, deeply buried, or multi-domain complexes.

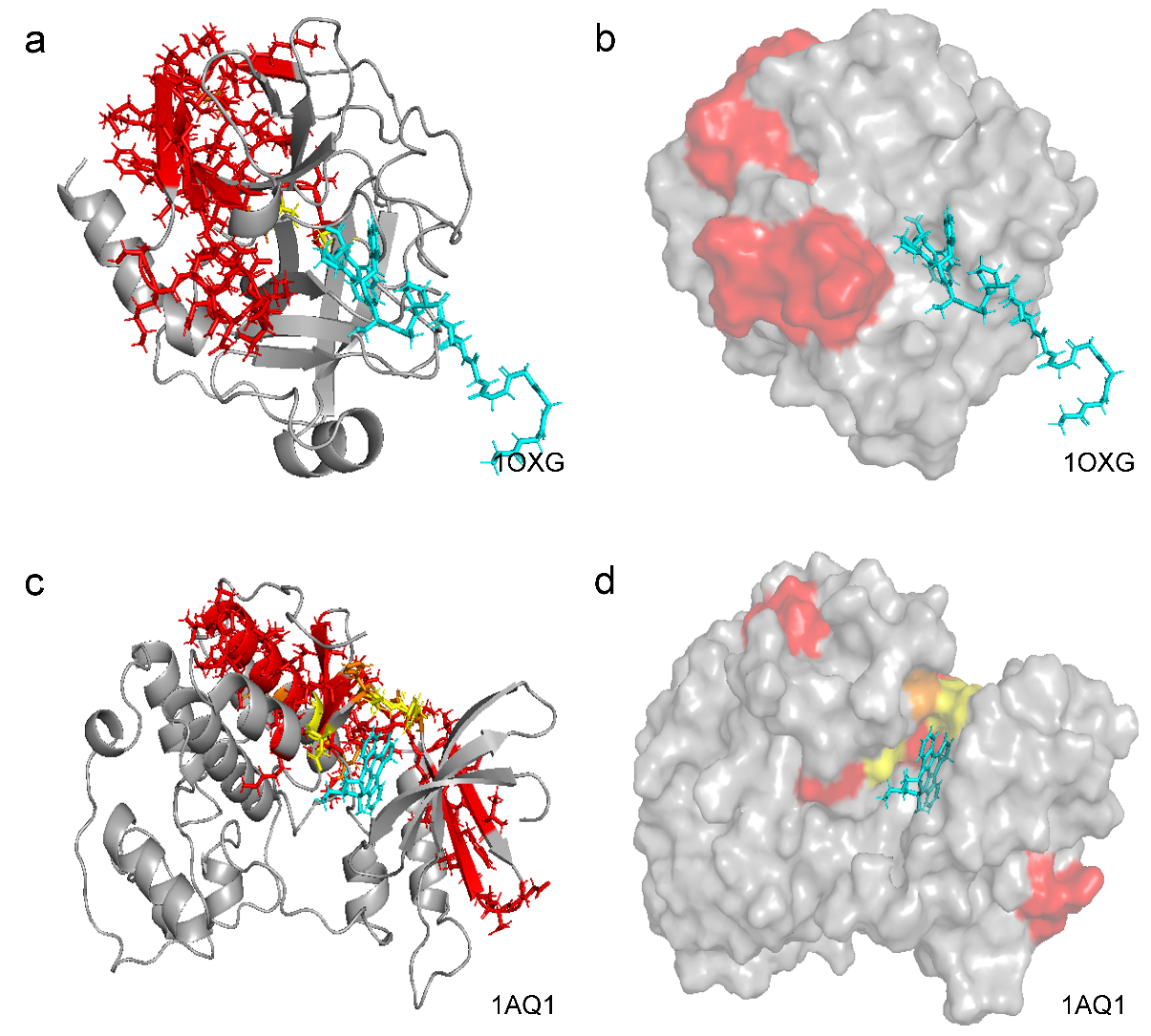

Supplementary Fig. S10 | Structural validation of binding site localization for 1OXG and 1AQ1. 3D visualizations of 1OXG (a, b) and 1AQ1 (c, d). Panels (a) and (c) display ribbon representations, while panels (b) and (d) show molecular surface representations to demonstrate ligand accommodation. Residues are colored by attention frequency: red (>75%), orange (50–75%), and yellow (25–50%). Although these targets show lower pixel-level precision compared to 5HLS, the multi-colored clusters in the surface views (b, d) still spatially converge near the cyan ligand. The broad distribution of yellow/orange residues likely reflects the challenge of resolving multi-domain binding interfaces using only 1D sequence data.

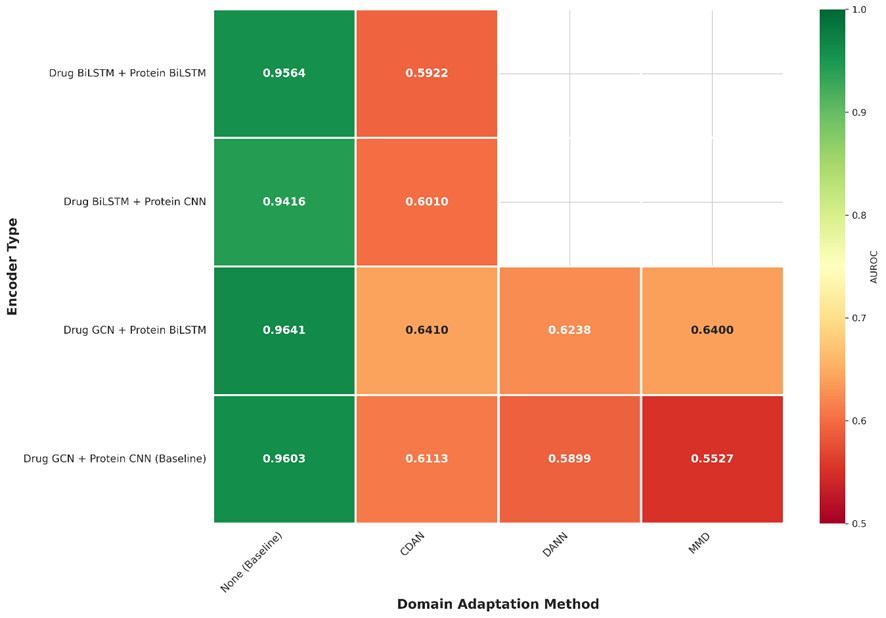

Supplementary Fig. S11 | Impact of domain adaptation on predictive performance. A heatmap matrix comparing AUROC scores across different encoder architectures with and without domain adaptation methods, including Conditional Adversarial Domain Adaptation (CDAN), Domain-Adversarial Neural Networks (DANN), and Maximum Mean Discrepancy (MMD). The color gradient represents the AUROC value, highlighting that the inclusion of CDAN consistently degrades performance compared to the baseline model without domain adaptation.

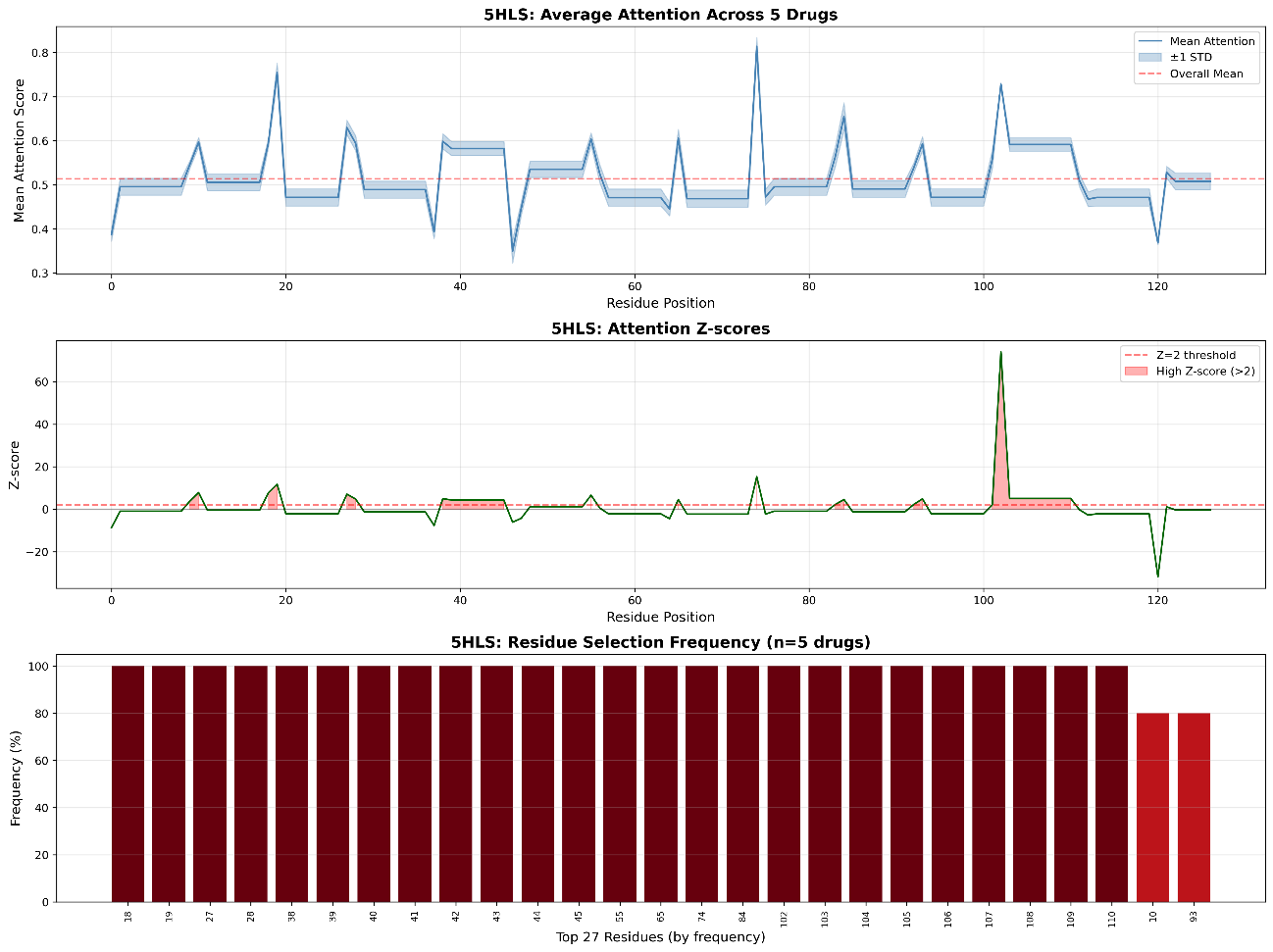

Supplementary Fig. S12 | Learned attention profile for 5HLS using the CNN-based baseline. The visualization displays the mean attention score (top) and standardized Z-scores (middle) across the amino acid sequence, alongside residue selection frequency (bottom) for the target 5HLS (Bromodomain). In contrast to the continuous attention motifs generated by our GCN-BiLSTM framework, the CNN model produces a highly discontinuous profile characterized by sharp, isolated spikes (e.g., the extreme Z-score peak near residue 100). This fragmented sequence-level signal indicates that the local receptive fields of the CNN encoder fail to integrate the global sequence context, directly leading to the scattered and inaccurate spatial predictions discussed in the structural analysis.

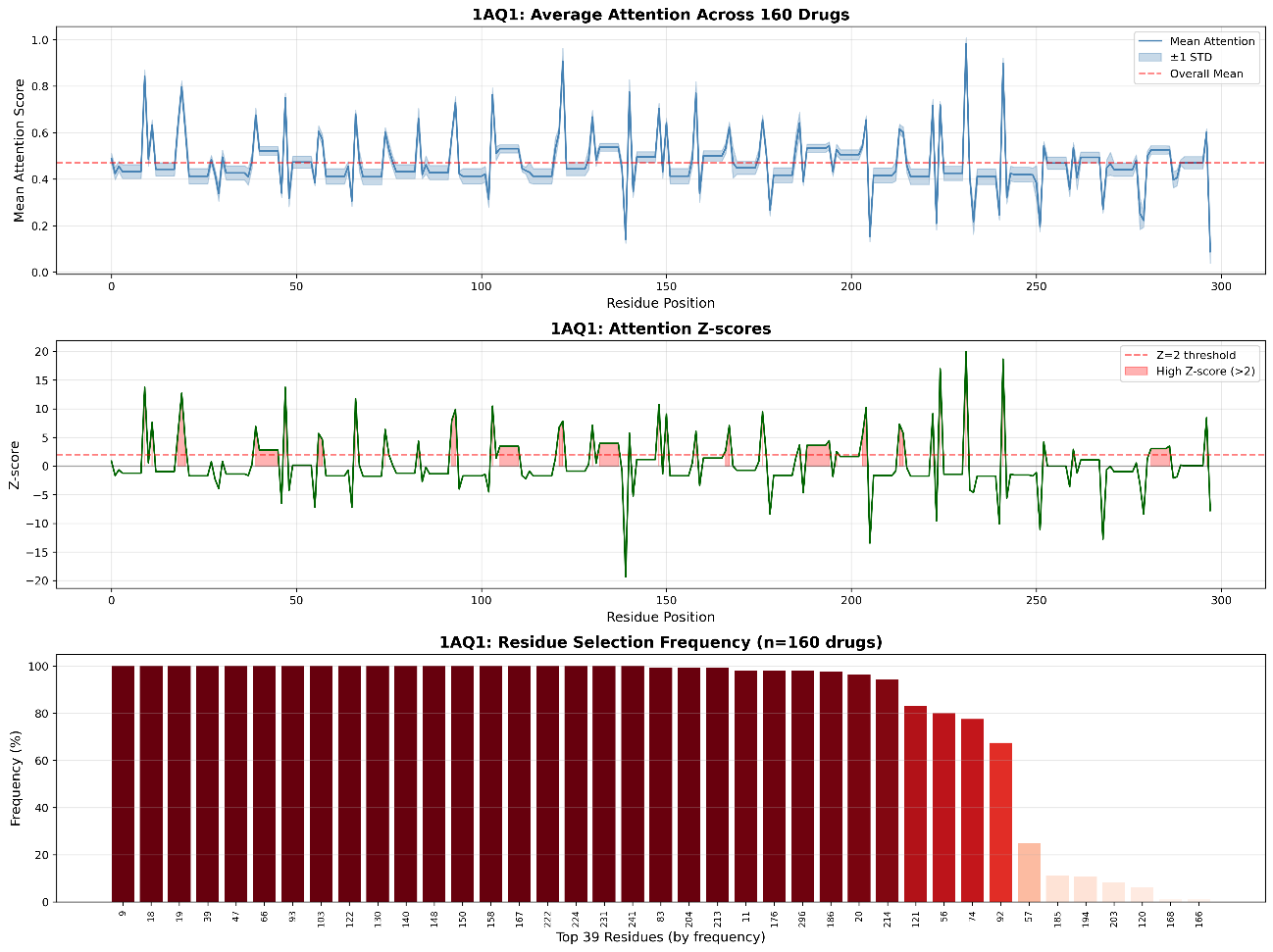

Supplementary Fig. S13 | Learned attention profile for 1AQ1 using the CNN-based baseline. The visualization displays the mean attention score (top) and standardized Z-scores (middle) across the amino acid sequence, alongside residue selection frequency (bottom) for the target 1AQ1. In contrast to the BiLSTM-derived profile (Supplementary Fig. S1), which exhibits distinct, continuous peaks corresponding to structural motifs, the CNN model generates high-frequency, discontinuous spikes. This fragmentation indicates that the local receptive fields of the CNN encoder fail to capture the sequential dependencies required to identify coherent binding regions, directly resulting in the dispersed structural prediction shown in Supplementary Fig. S22 c, d.

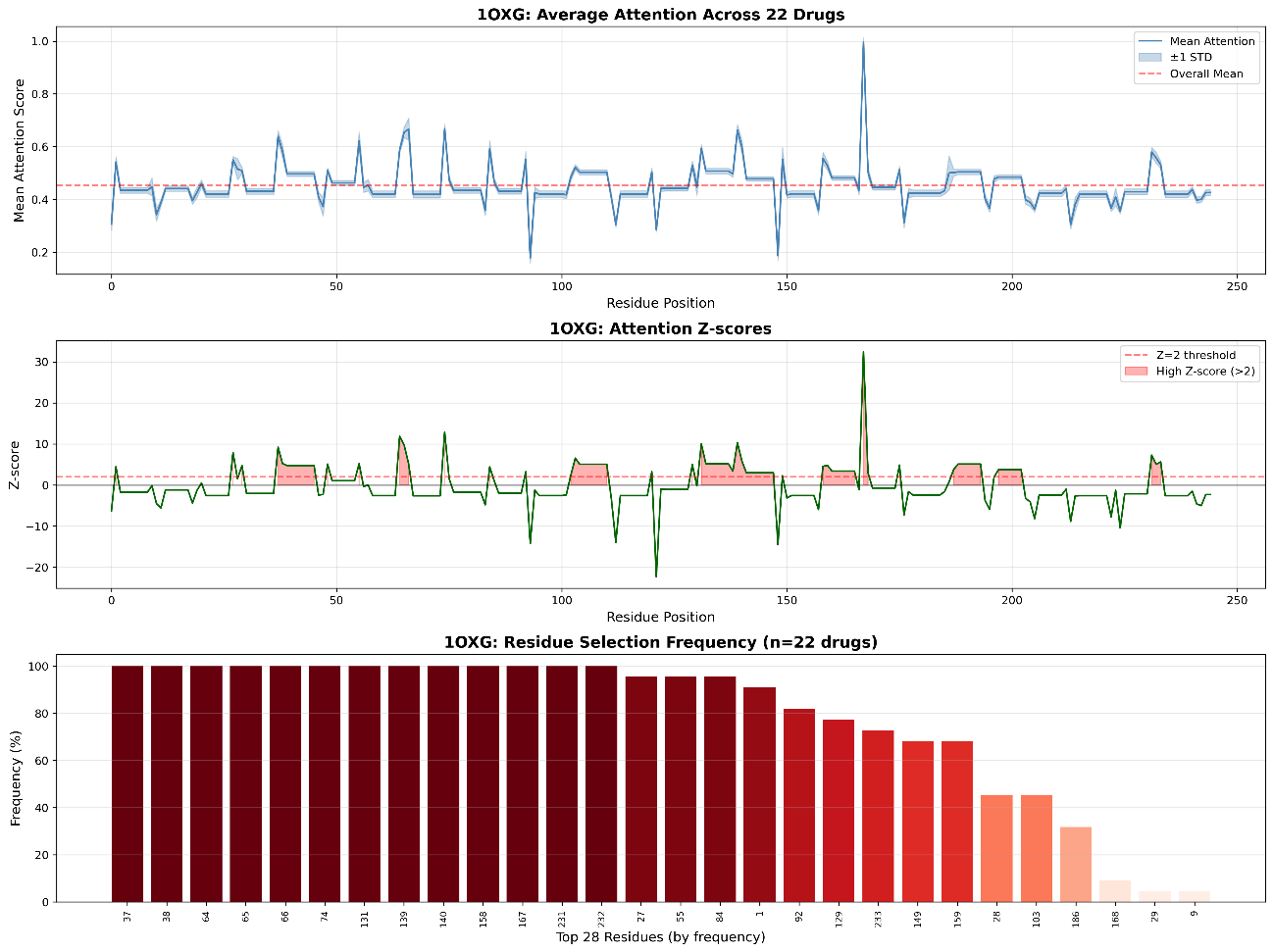

Supplementary Fig. S14 | Learned attention profile for 1OXG using the CNN-based baseline. Analogous to Supplementary Fig. S13, this figure displays the attention distribution for the target 1OXG using the CNN encoder. Compared to the BiLSTM baseline (Supplementary Fig. S2), which shows a clear accumulation of attention weights in specific domains, the CNN encoder produces a noisy signal characterized by scattered peaks with no clear topological structure. This “spiky” distribution lacks the contextual continuity observed in the BiLSTM profile, leading to the spatial dispersion observed in the 3D structural mapping (Supplementary Fig. S22 a, b).

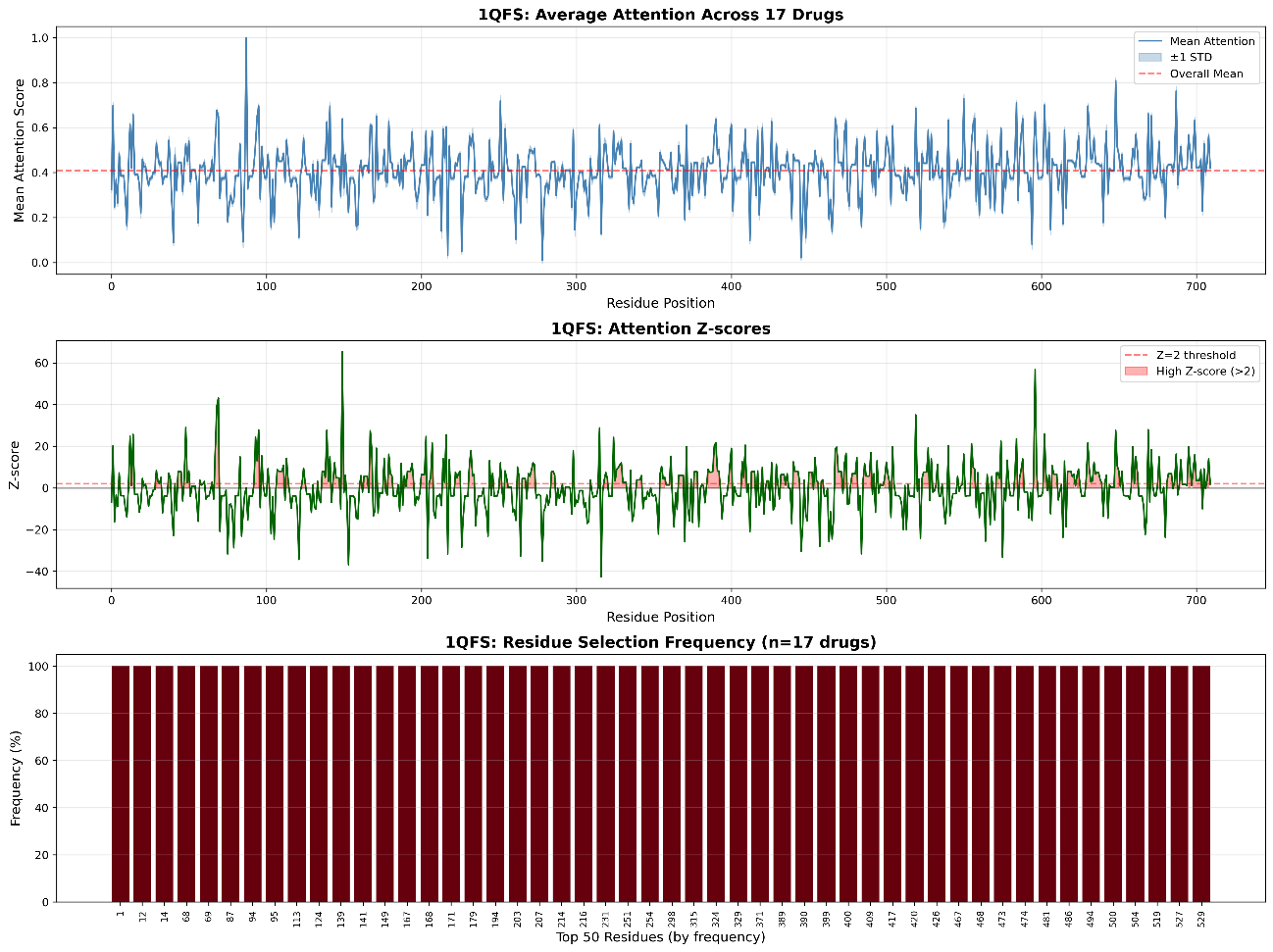

Supplementary Fig. S15 | Learned attention profile for 1QFS using the CNN-based baseline. The visualization displays the mean attention score (top) and standardized Z-scores (middle) across the amino acid sequence, alongside residue selection frequency (bottom) for the target 1QFS. This target represents a challenging multi-domain structure where the BiLSTM model previously exhibited limitations (Supplementary Fig. S3). However, the CNN encoder produces an even more erratic signal, characterized by dense, high-frequency noise across the entire sequence length. Unlike the BiLSTM model, which attempts to define (albeit incorrectly) specific regions, the CNN model fails to distinguish any meaningful structural motifs, leading to the highly scattered structural prediction shown in Supplementary Fig. S21 c, d.

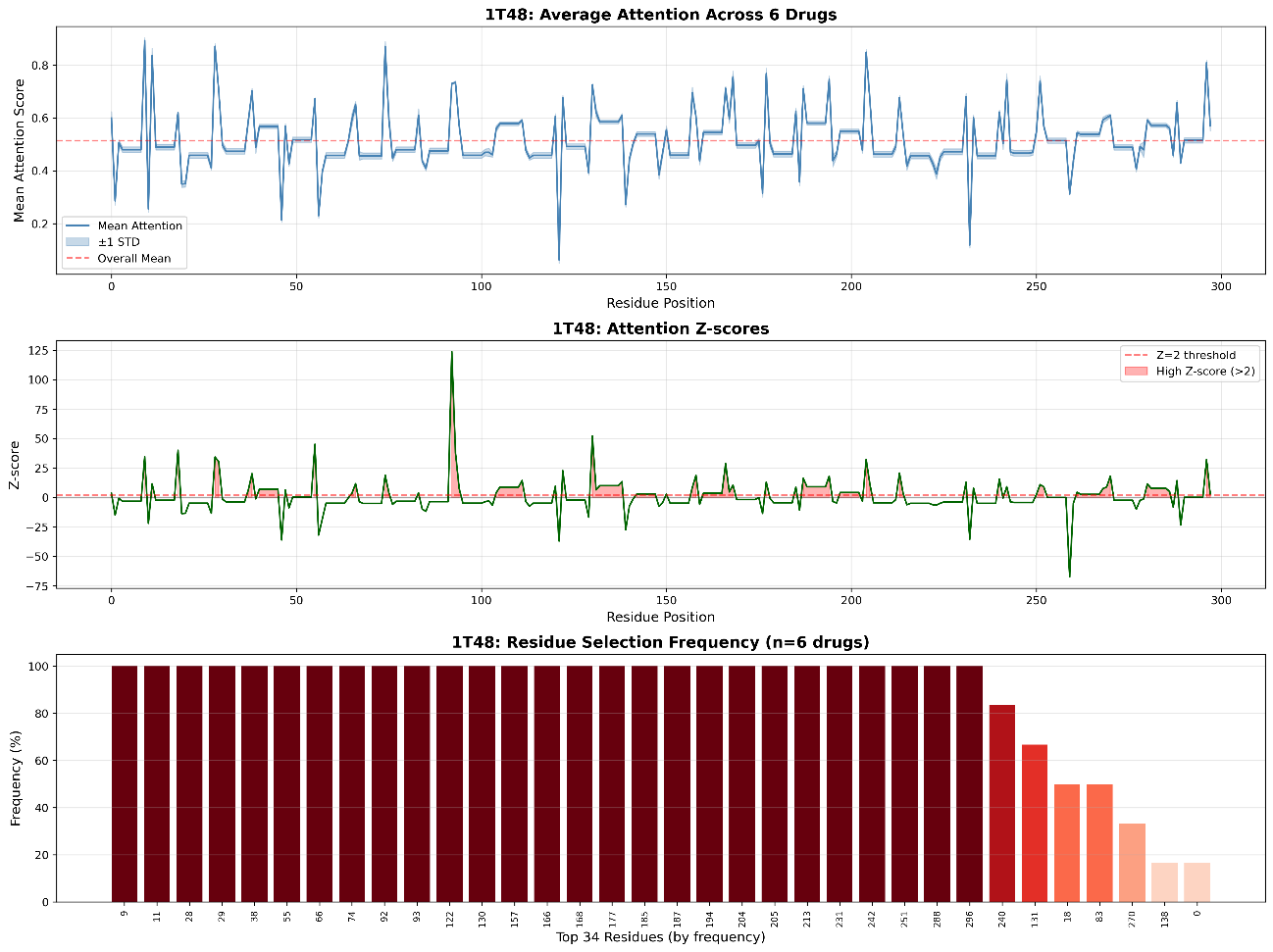

Supplementary Fig. S16 | Learned attention profile for 1T48 using the CNN-based baseline. The visualization displays the mean attention score (top) and standardized Z-scores (middle) across the amino acid sequence, alongside residue selection frequency (bottom) for the target 1T48 using the CNN encoder. In direct comparison with the BiLSTM-derived profile (Supplementary Fig. S4), which exhibits clear, wave-like regions of importance, the CNN model generates a fragmented and noisy profile. The lack of sequential continuity indicates that the local convolution filters struggle to integrate context over this sequence, resulting in isolated spikes that do not correspond to the continuous binding interface, as evidenced by the diffuse structural prediction in Supplementary Fig. S21 a, b.

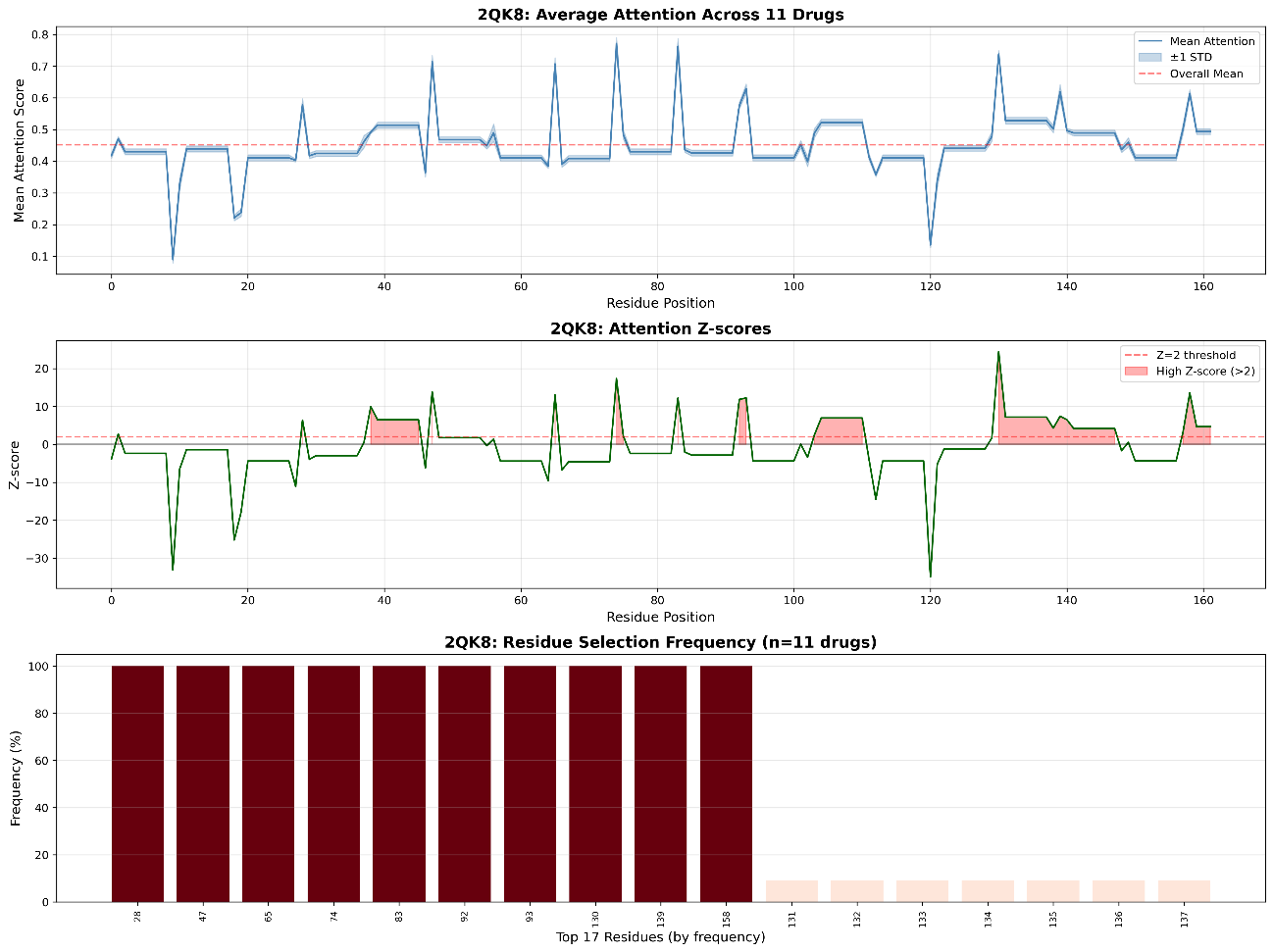

Supplementary Fig. S17 | Learned attention profile for 2QK8 using the CNN-based baseline. The visualization displays the mean attention score (top) and standardized Z-scores (middle) across the amino acid sequence, alongside residue selection frequency (bottom) for the target 2QK8 using the CNN encoder. In contrast to the BiLSTM-derived profile (Supplementary Fig. S5), which revealed distinct high-attention regions corresponding to structural anchors, the CNN model generates a chaotic distribution characterized by dense, high-frequency fluctuations. This lack of signal clarity suggests that the CNN encoder fails to filter out background noise or identify the dominant sequence motifs required for ligand binding, directly contributing to the dispersed structural results shown in Supplementary Fig. S20 c, d.

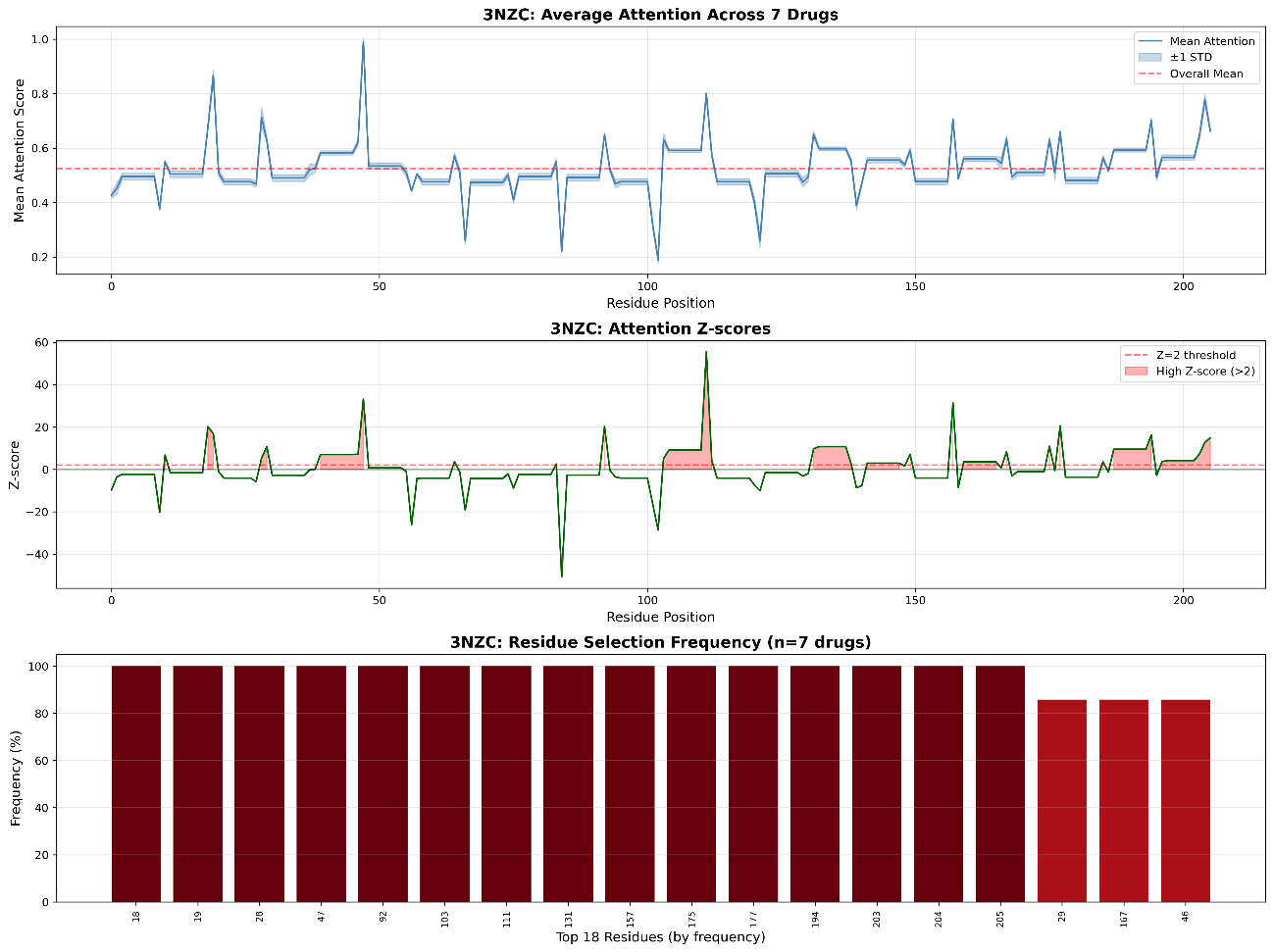

Supplementary Fig. S18 | Learned attention profile for 3NZC using the CNN-based baseline. The visualization displays the mean attention score (top) and standardized Z-scores (middle) across the amino acid sequence, alongside residue selection frequency (bottom) for the target 3NZC (Bromodomain). This profile provides the sequence-level explanation for the structural results presented in the main text. In direct contrast to the BiLSTM-derived profile (Supplementary Fig. S6), which displayed smooth, wave-like attention motifs corresponding to the binding domain, the CNN model generates high-frequency, discontinuous spikes. This lack of sequential continuity highlights the limitation of fixed local receptive fields. Crucially, this fragmented sequence profile provides the mechanistic basis for the model's failure in the structural domain, directly resulting in dispersed residue predictions where the CNN baseline fails to spatially converge around the ligand.

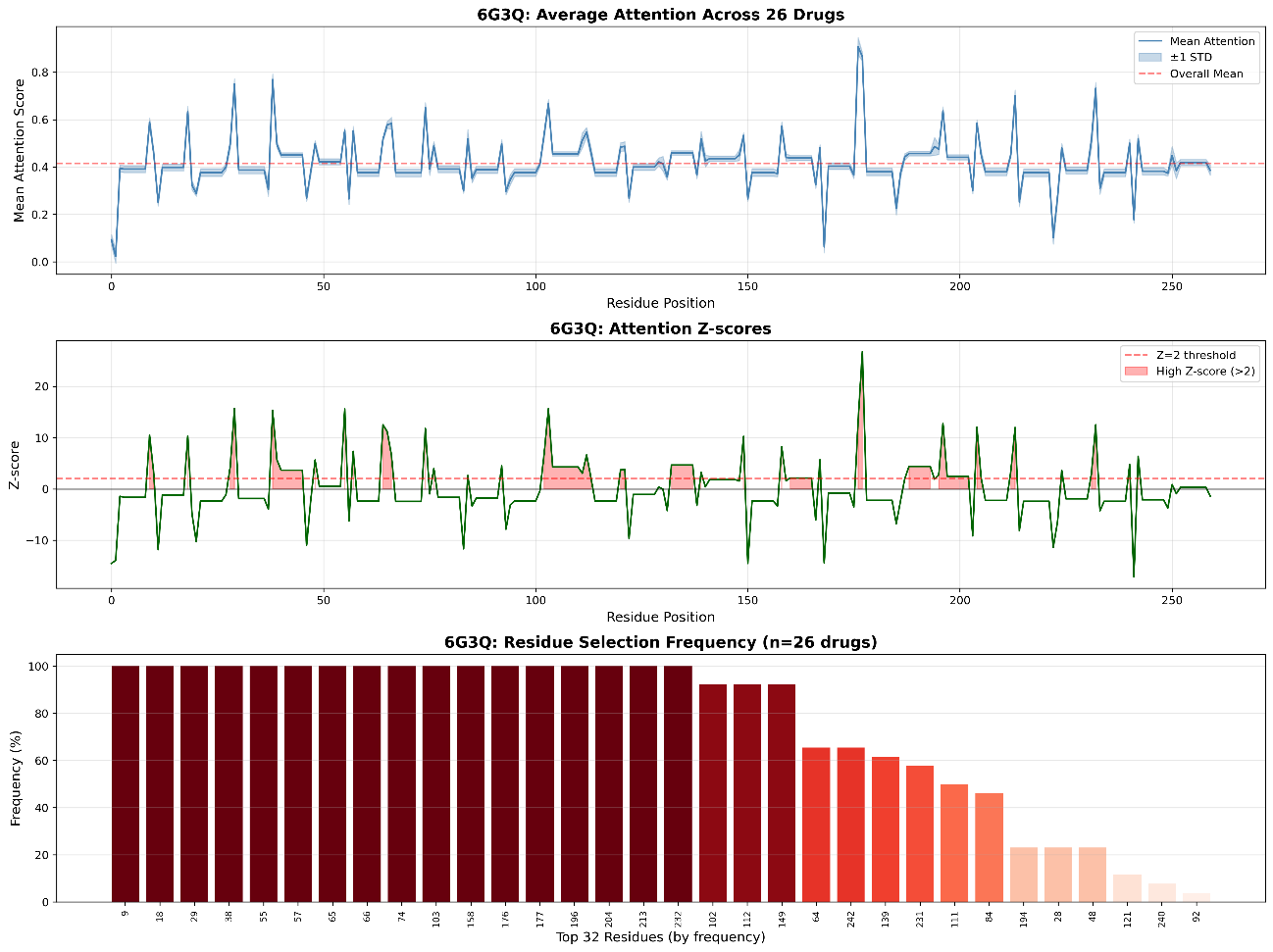

Supplementary Fig. S19 | Learned attention profile for 6G3Q using the CNN-based baseline. The visualization displays the mean attention score (top) and standardized Z-scores (middle) across the amino acid sequence, alongside residue selection frequency (bottom) for the target 6G3Q using the CNN encoder. Comparison with the BiLSTM baseline (Supplementary Fig. S7) highlights a fundamental deficit in the CNN architecture. While the BiLSTM model produced a clean, peak-driven profile indicative of a learned global context, the CNN model yields a noisy, fragmented profile. The inability of the local convolution filters to integrate long-range dependencies results in a series of disconnected spikes that fail to represent a coherent binding domain, leading to the diffuse spatial distribution observed in Supplementary Fig. S20 a, b.

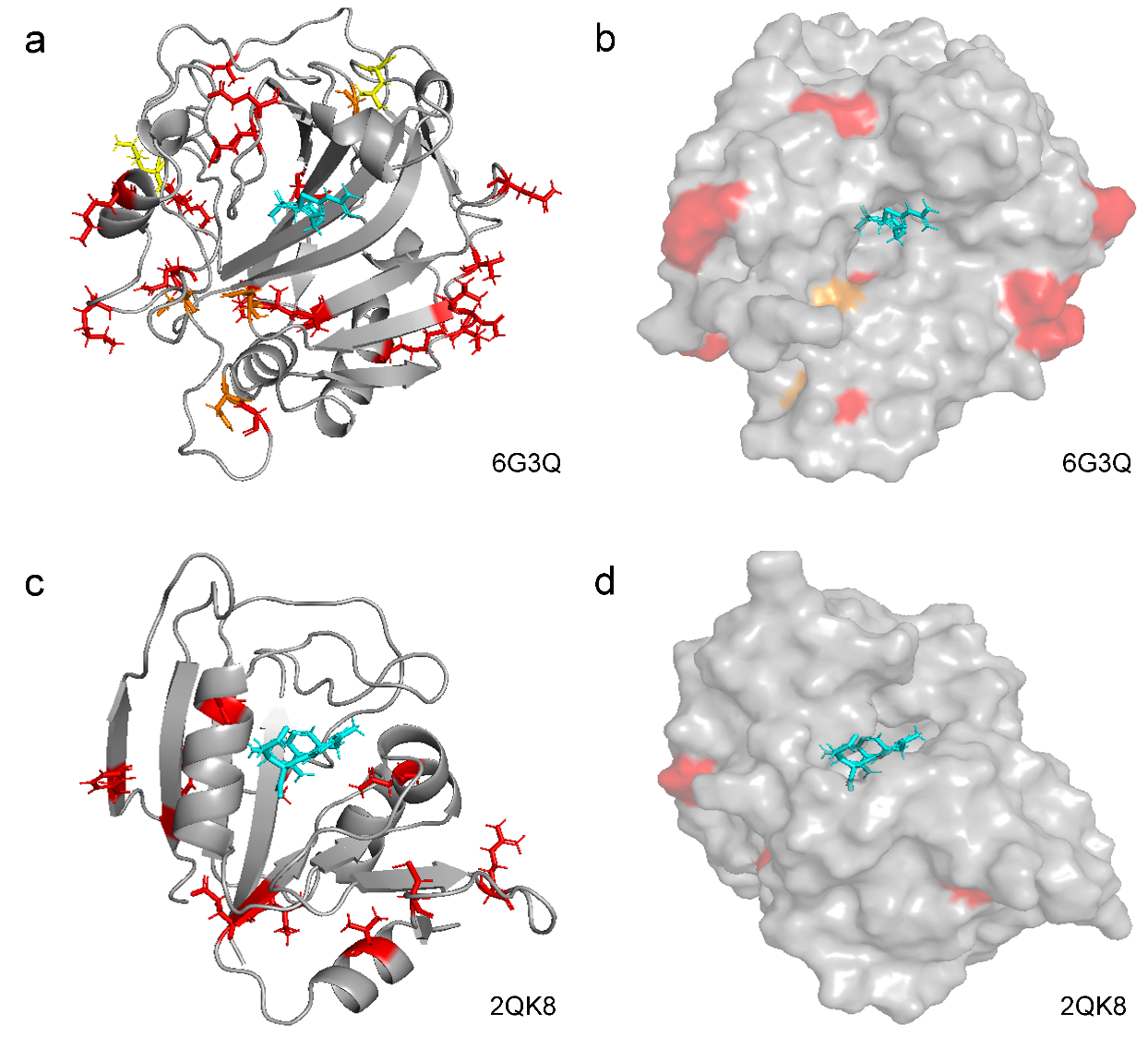

Supplementary Fig. S20 | Structural validation of binding site localization for 6G3Q and 2QK8 using the CNN-based baseline. 3D visualizations of the attention-derived binding sites for targets 6G3Q (a, b) and 2QK8 (c, d) using the DrugBAN (CNN) encoder. Panels (a) and (c) display ribbon representations; panels (b) and (d) show molecular surfaces. Residues are colored by attention frequency: red (>75%), orange (50–75%), and yellow (25–50%). For 6G3Q (a, b), unlike the BiLSTM results (Supplementary Fig. S8 a, b), which supported a “functional hierarchy” where anchor residues clustered centrally, the CNN model predicts a diffuse cloud of residues scattered across the protein surface without structural logic. Similarly, for 2QK8 (c, d), the CNN baseline fails to localize the binding pocket. The high-confidence residues are dispersed distally from the ligand (cyan), exhibiting none of the spatial convergence observed in the BiLSTM model (Supplementary Fig. S8 c, d). This comparison serves as robust evidence that the BiLSTM encoder’s ability to capture global sequence context is a prerequisite for generating interpretable, structure-aware predictions in the absence of 3D supervision.

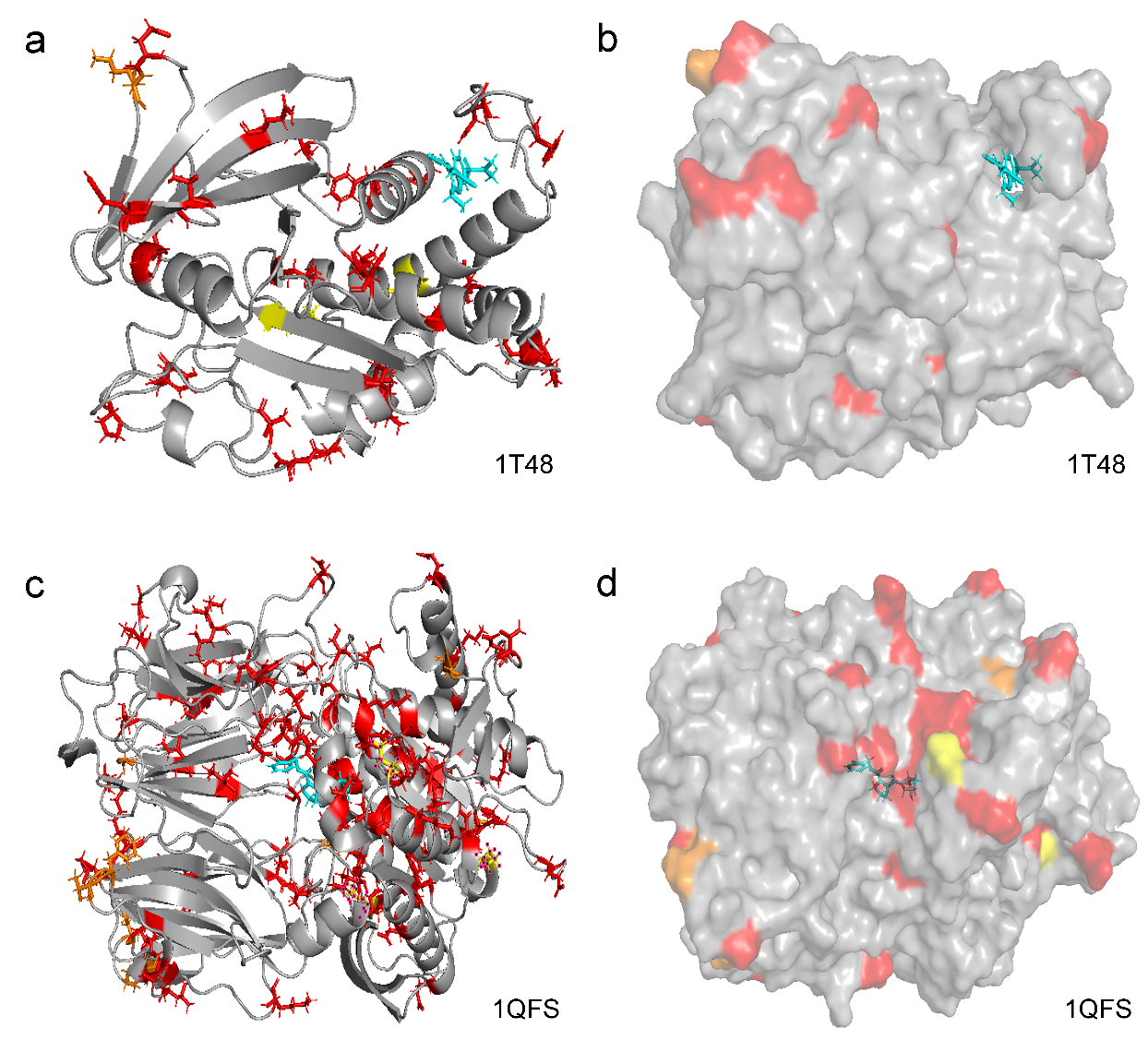

Supplementary Fig. S21 | Structural validation of binding site localization for 1T48 and 1QFS using the CNN-based baseline. 3D visualizations of the attention-derived binding sites for targets 1T48 (a, b) and 1QFS (c, d) using the DrugBAN (CNN) encoder. Panels (a) and (c) display ribbon representations; panels (b) and (d) show molecular surfaces. Residues are colored by attention frequency: red (>75%), orange (50–75%), and yellow (25–50%). For 1T48 (a, b), the contrast is pronounced. While the BiLSTM model identified the general binding region (Supplementary Fig. S9 a, b), the CNN model produces a diffuse signal with high-frequency residues scattered distally from the ligand (cyan). This further demonstrates that the local receptive fields of the CNN encoder are insufficient for localizing binding pockets in targets where global context is critical. For 1QFS (c, d), which was identified as a challenging boundary condition for the BiLSTM framework (Supplementary Fig. S9 c, d), the CNN baseline similarly fails to identify the binding pocket. However, it exhibits a distinct topological error: the high-confidence residues are randomly dispersed across multiple domains, reflecting the model's inability to handle large, complex sequences.

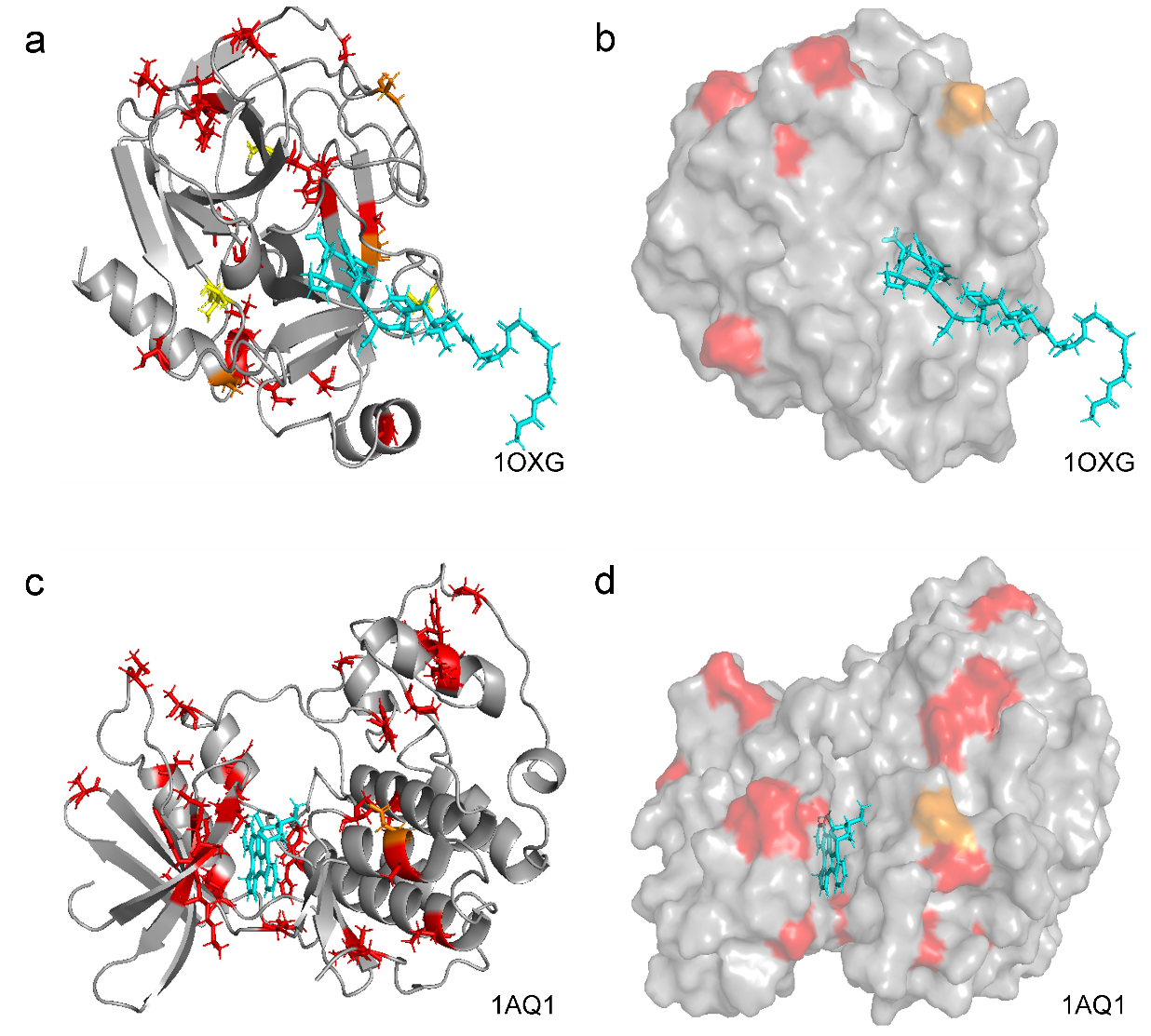

Supplementary Fig. S22 | Structural validation of binding site localization for 1OXG and 1AQ1 using the CNN-based baseline. 3D visualizations of the attention-derived binding sites for targets 1OXG (a, b) and 1AQ1 (c, d) using the DrugBAN (CNN) encoder. Panels (a) and (c) display ribbon representations, while panels (b) and (d) show molecular surface representations to demonstrate spatial distribution. Residues are colored by attention frequency: red (>75%), orange (50–75%), and yellow (25–50%). For 1OXG (a, b), the CNN prediction fails to form a consolidated binding interface, standing in sharp contrast to the coherent pocket identified by the BiLSTM model (Supplementary Fig. S10 a, b). The high-frequency residues (red) are scattered randomly across the protein surface. Similarly, for 1AQ1 (c, d), unlike the BiLSTM model which successfully localized a spatially converged pocket even for this challenging target (Supplementary Fig. S10 c, d), the CNN model yields a fragmented distribution where high-confidence residues are dispersed without structural logic. These structural failures are direct consequences of the discontinuous attention profiles shown in Supplementary Figs. S14 (for 1OXG) and S13 (for 1AQ1), confirming that the CNN encoder lacks the global contextual modeling capabilities necessary for accurate binding site identification in complex folds.

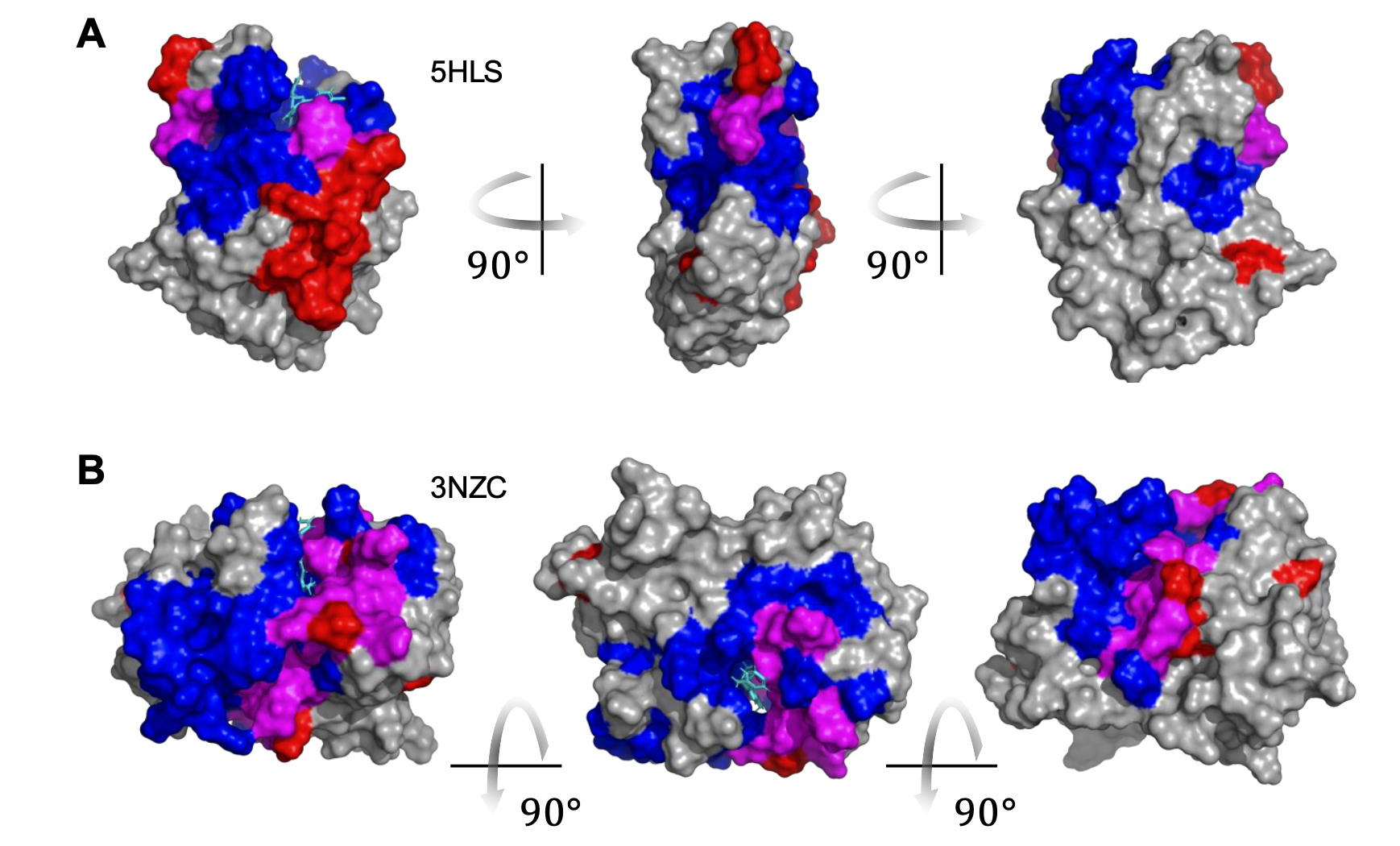

Supplementary Fig. S23 | Synergistic visualization of geometric (fpocket) and semantic (GCN-BiLSTM) predictions. Surface representations of predicted binding sites for (a) 5HLS and (b) 3NZC across three orthogonal rotations ($90^{\circ}$). This visualization facilitates a direct comparison between sequence-based attention (semantic) and geometric cavity detection. Color coding highlights the spatial consensus and divergence between methods: regions uniquely identified by GCN-BiLSTM attention are shown in red, while those uniquely identified by fpocket geometric analysis are in blue. The magenta regions represent the intersection where both semantic attention and geometric voids overlap. The concentration of magenta patches near the crystallized ligands (cyan) demonstrates the “functional filtering” effect, where BiLSTM attention effectively narrows down geometric candidates to the biologically active orthosteric site.
